# The role of IgE patterns and their link to the gut microbiome in allergic sensitization

**DOI:** 10.64898/2026.03.25.714318

**Authors:** Oleg Vlasovets, Marie Standl, Lisa Maier, Stefanie Gilles, Harald Grallert, Claudia Traidl-Hoffmann, Annette Peters, Christian L. Müller

## Abstract

Allergic diseases are heterogeneous conditions shaped by immune pathways and environmental influences, including the gut microbiome. Using cross-sectional data from 508 adults in the KORA FF4 cohort (275 with IgE sensitization, 233 without), we provide a multimodal statistical analysis of deep IgE profiles and concomitant gut microbial amplicon sequencing variant (ASV) data. We identified three latent allergy components in the cohort’s IgE profiles that reflects food, pollen, and house dust mite markers and enables interpretable stratification of the cohort. Contrary to prior studies, microbial diversity did not differ between sensitized and non-sensitized individuals across all cohort strata. Differential abundance analysis identified 61 ASVs, with enrichment in *Bacteroidaceae*, *Oscillospiraceae*, and *Veillonellaceae* families, and depletion in the *Lachnospiraceae* family. Microbial network analysis further identified altered family-level associations in pollen- and food-sensitized individuals. Taxon set enrichment highlighted folic acid- and vitamin A-producing taxa, with consistent signals from *Prevotella copri* (depleted) and *Bacteroides massiliensis* (enriched). Together, our analysis points toward specific microbial families and metabolic groups as correlates of IgE sensitization in an adult population.

**HIGHLIGHTS:**

- IgE sensitization clusters reflect allergen sources and cross-reactive proteins
- Three latent allergy components capture the IgE profile structure of the entire cohort
- Distinct IgE patterns are linked to changes in taxa compositions and associations
- Folic acid– and vitamin A–producing taxa are enriched in IgE-sensitized individuals

**GRAPHICAL ABSTRACT:** 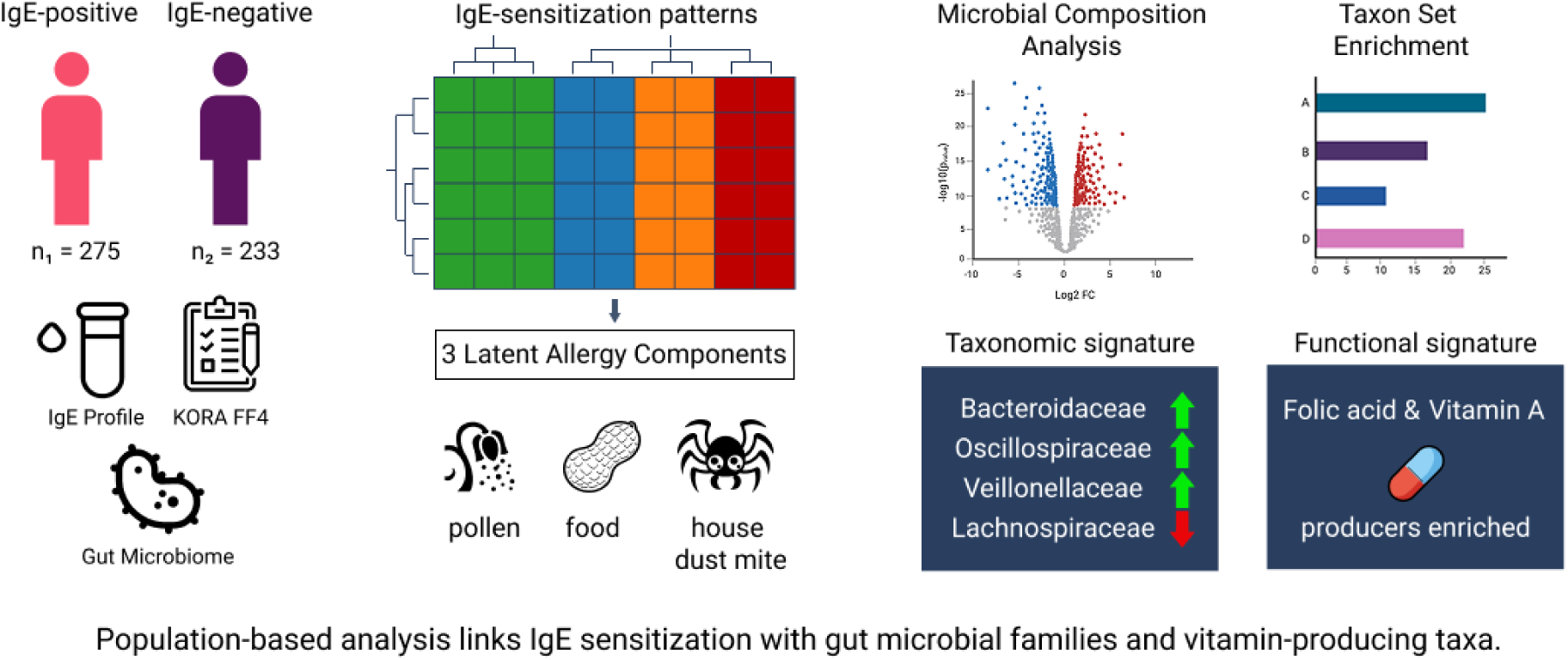

## 1 INTRODUCTION

Allergic diseases are highly prevalent, affecting more than 400 million individuals worldwide and ranging from common conditions such as allergic rhinitis to severe conditions such as anaphylaxis [1]. These disorders represent chronic inflammatory responses to harmless environmental substances. Among the diverse immunological mechanisms, Immunoglobulin E (IgE)–mediated allergies correspond to Type I hypersensitivity and remain central to both research and clinical practice. While the effector mechanisms of IgE-mediated allergies are well described [2], the mechanisms driving sensitization as well as patterns of co-sensitization across individuals remain poorly understood [3].

To gain a more quantitative understanding of the relationship between the underlying molecular causes and a patient’s allergy phenotype, researchers and clinicians increasingly rely on advanced molecular allergy testing technologies, such as, e.g., Immuno Solid-phase Allergen Chips (ISACs) [4], which provide cost-effective measurements of deep multiplexed IgE profiles at cohort scale. Linking these IgE profiles with concomitant patient health or ‘omics’ data holds the promise to identify patterns of cross-reactivity [5], allergic disease endotypes [6], and ultimately, hypotheses for biological mechanisms. For instance, clustered IgE profiles have already been shown to be linked to asthma and duration of allergy in a retrospective clinical cohort in Vienna [7]. Similarly, statistical analysis of the cross-sectional population-based health survey cohort for Luxembourg [8](EHES-LUX) together with IgE profiles have revealed strong associations between a population sub-group with high IgE burden and physician-diagnosed asthma as well as nasal, eye, and food allergy [9].

Among the many factors that contribute to allergy development and progression, including an individual’s genetics, living conditions, and environmental exposure, microbiota have been implicated as a critical component for shaping immune tolerance and allergic predisposition [10–12]. For example, alterations in the intestinal microbiome, as measured by 16S rRNA sequencing, have been associated with asthma and allergic rhinitis in a small pediatric cohort [13]. Moreover, coevolution of humans and their gut microbiota suggests that microbes may contribute to IgE-mediated food allergy through immune dysregulation and impaired gut barrier function [14]. Recent evidence indicates that the intestinal mucus layer represents a critical but overlooked component in allergic sensitization [15]. This glycoprotein-rich barrier serves not only as a physical separation between luminal antigens and the epithelium, but also as a platform for immune regulation and oral tolerance induction [15]. Importantly, the integrity of this barrier depends on a delicate balance between mucin-degrading bacteria, such as *Bacteroides* species and *Akkermansia muciniphila*, and protective, short-chain fatty acid (SCFA) producing taxa, particularly members of Lachnospiraceae [16, 17]. Indeed, in a seminal study, Goldberg et al. [18] reported significantly different gut microbial 16S rRNA sequencing-based signatures in a cohort of 233 patients (age *>* 48 months) with food allergy (either confirmed by a physician or by oral food challenge), compared to age-matched non-allergic controls. Both microbial alpha and beta diversity were significantly different, with *Prevotella copri* (*P. copri*) being the most overrepresented species in non-allergic controls. Moreover, metabolomics-derived SCFAs levels were significantly higher in the non-allergic control group compared to the food allergy group, suggesting a strong relationship between IgE status and gut microbial composition and its associated metabolic profile.

To further advance our understanding of the relationship between IgE sensitization patterns and the gut microbiome, we present a multimodal large-scale integrative analysis of deep allergen-specific IgE profiles and gut microbiome amplicon sequencing data from 508 adults in the population-based KORA FF4 cohort. Our IgE profiles, comprising 112 allergen-specific markers, cover all major allergens, including food, pollen, animal dander, and host dust mite markers, thus allowing for a fine-grained characterization and stratification (sub-grouping) of the cohort and a more general analysis of IgE sensitization-microbe relationships.

We show that previously observed patterns of microbial diversity changes [18] in allergy-sensitized individuals cannot be reproduced in the present adult cohort, and that microbial compositions are only weakly predictive of IgE sensitization patterns, both cohort-wide and with respect to particular sub-groups (strata). We can, however, identify specific microbial taxa in the families *Bacteroidaceae*, *Oscillospiraceae*, *Veillonellaceae*, and *Lachnospiraceae* that are associated with certain cohort sub-groups. Furthermore, we show altered microbial family-family association patterns between IgE sensitized and non-sensitized groups. In the absence of metabolomics data for the present cohort, we supplement our analysis with an in-depth functional taxon set enrichment analysis, revealing a further link between IgE sensitization and Vitamin A and folic acid producers. Taken together, our results produce several testable hypothesis in form of microbial taxa, families, and metabolite producers that may play a pivotal role in IgE sensitization in adults.

## 2 RESULTS

Our analysis proceeds from patterns of IgE sensitization to their microbial correlates. We first define co-sensitization clusters and derive latent allergy components, then test whether these immunological patterns are reflected in global microbial diversity or in shifts of specific taxa abundances. We next assess the ability of microbial profiles to predict IgE sensitization, evaluate differences in microbial network structure, and finally apply taxon set enrichment analysis to link microbial functional potential to IgE status.

### IgE sensitization is linked to a higher prevalence of asthma, dermatitis, and hay fever in the KORA FF4 cohort

We analyzed 508 individuals from the KORA FF4 cohort, including 275 with IgE sensitization and 233 without. The study design is shown in Figure S1, and descriptive statistics are summarized in Table 1. IgE sensitization was defined as the presence of an allergen-specific IgE for at least one tested allergen.

**Table 1.**
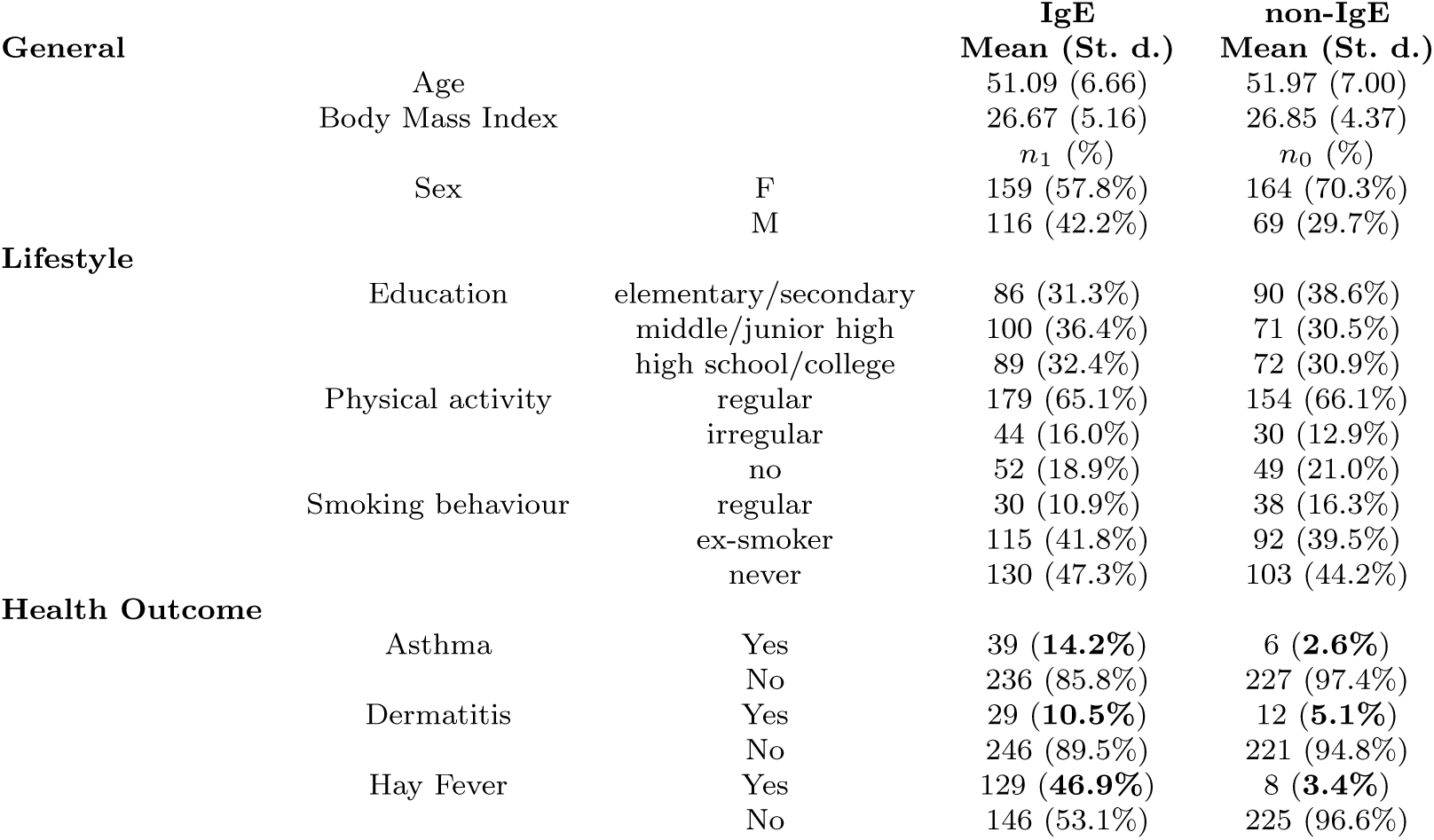
Summary statistics of the KORA study cohort including demographic, lifestyle, and health outcome variables across IgE-positive and IgE-negative individuals.

Demographic characteristics such as age and BMI were comparable between sensitized and non-sensitized groups. A higher proportion of females was observed in the non-sensitized group (70.3%) compared to the sensitized group (57.8%). Lifestyle factors, including education, physical activity, and smoking, showed little variation between groups. By contrast, sensitized individuals exhibited markedly higher prevalence of clinical allergic conditions: asthma (14.2% versus 2.6%), dermatitis (10.5% versus 5.1%), and hay fever (46.9% versus 3.4%). These differences confirm that IgE sensitization is associated with increased burden of allergic disease in the study population.

### Three latent allergy components capture major IgE-sensitization patterns

We next sought to identify major IgE-sensitization patterns of the 112 allergen-specific markers across the 275 sensitized individuals. We first binarized the measured IgE response values, resulting in a 112×275 presence-absence matrix *Q* (Figure 1A top panel). We applied hierarchical clustering to the binarized IgE matrix using pairwise Jaccard distances and complete linkage separately for rows (IgEs) and columns (individuals). Clustering of the individuals identified roughly nine subpopulations (groups A-I in Figure 1A top panel) with distinct IgE patterns. Clustering of IgE patterns across the cohort revealed multiple clusters with three dominant groupings, characterized by food-related IgEs (marked in red), pollen-related IgEs (marked in green), and house dust mite (HDM)–related IgEs.

**Fig. 1.**
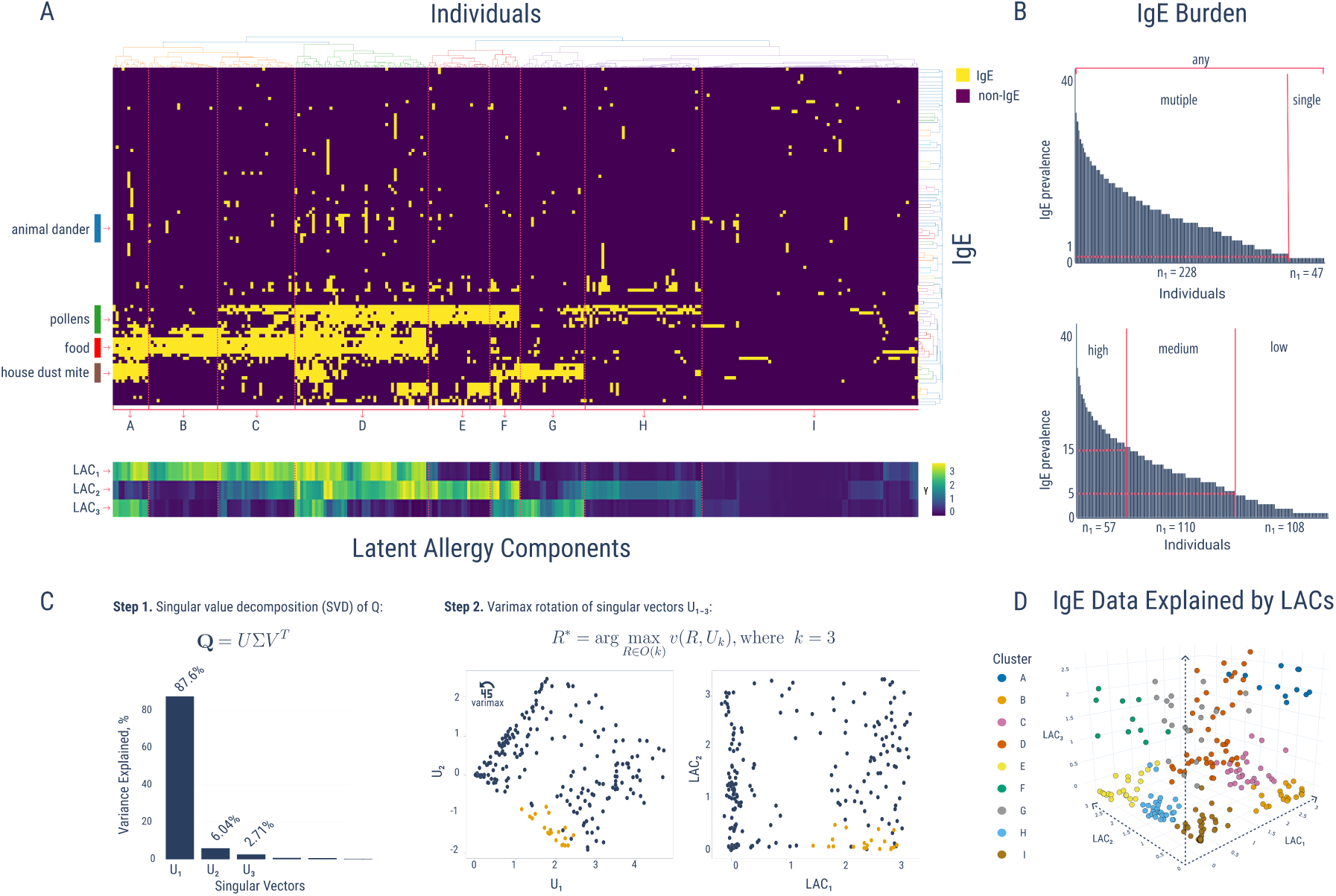
IgE sensitization clusters define latent allergy components (A) Hierarchical clustering of binary IgE profiles from 275 sensitized individuals reveals nine clusters (A–I). Latent Allergy Components (LACs), derived from low-rank factorization of IgE data, are shown below the heatmap with component scores ranging from 0 to 3. (B) Partitioning of individuals by IgE Burden Score, defined as the number of detected allergen-specific IgEs. (C) Scree plot of variance explained by each LAC, with scatter plot of IgEs in rotated LAC space highlighting improved interpretability.(D) Three-dimensional visualization of the LAC space showing distinct spatial separation of individuals, colored by cluster identity.

Indeed, when performing a singular value decomposition (SVD) of the *Q*-matrix, we observed that more than 96% of variance in the IgE data can be explained by the first three SVD components (Figure 1C, Step 1). Projection of the 275 samples onto the three components exhibits a rotated orthogonal patterning that we turned into interpretable orthogonal allergy components via varimax rotation [19] (Figure 1C, Step 2). We refer to these three major axis of variations as latent allergy components (LACs) of the cohort. Visualization of the LACs (Figure 1A lower panel) showed that LAC1 largely reflects food-related IgEs (marked in red), LAC2 pollen-related IgEs (marked in green), and LAC3 house dust mite (HDM)–related IgEs (marked in dark red). Joint analysis of the clustered groups and the latent allergy components confirmed strong agreement and allows interpretability of the different sub-populations. Figure 1D shows how the nine different sub-groups populate distinct axes and corners of the three-dimensional latent allergy space. For instance, Cluster B has large loadings only in LAC1, thus representing participants carrying mostly food-related IgEs (colored in orange). Likewise, Cluster E has large loadings only in LAC2, representing participants with pollen-related IgEs (colored in yellow). Cluster G has large loadings only in LAC3, representing participants with house dust mite IgEs (marked in brown), while Cluster I exhibited consistently low values across all three components, thus representing a heterogeneous group with only a few disjoint IgE markers.

Table 2 describes the dominating IgE markers, demographic and clinical features of the data-driven IgE clusters (A–I). Birch (*Bet v 1*) dominated clusters B–D and appeared frequently in cluster I. Grass-pollen IgEs (*Phl p 1*, *Phl p 4*, *Phl p 5*, *Cyn d 1*) characterized clusters E, F, and H, while HDM components (*Der f 1*, *Der f 2*, *Der p 2*) were most common in clusters A and G. The heterogeneous Cluster I comprised *Ole e 1* (olive tree) and *Fel d 1* (cat) markers. Across all partitions, *Bet v 1* and *Phl p 1* were the most recurrent IgEs.

**Table 2.** Summary statistics of IgE clusters (A–I) including age, BMI, cluster size, and the most prevalent allergen-specific IgEs. Sex distribution is shown as percentage male/female, and education is reported as percentage with secondary (s), medium (m), and higher (h) education.

| Cluster | Top IgE Components | Size | Age (Mean, SD) | BMI (Mean, SD) | Sex (m/f %) | Education (s/m/h %) |
| --- | --- | --- | --- | --- | --- | --- |
| A | dust mite (*Der f 2*, *Der p 2*, *Der f 1*) | 12 | 51.17 (6.13) | 25.85 (2.76) | 50.0 / 50.0 | 25.0 / 50.0 / 25.0 |
| B | birch (*Bet v 1*), hazel (*Cor a 1*), apple (*Mal d 1*) | 24 | 53.88 (6.69) | 26.86 (5.81) | 37.5 / 62.5 | 20.8 / 41.7 / 37.5 |
| C | birch (*Bet v 1*), apple (*Mal d 1*), hazel (*Cor a 1*) | 26 | 51.81 (7.53) | 25.81 (4.94) | 23.1 / 76.9 | 11.5 / 50.0 / 38.5 |
| D | birch (*Bet v 1*), hazel (*Cor a 1*), apple (*Mal d 1*) | 45 | 48.80 (5.98) | 25.76 (3.99) | 48.9 / 51.1 | 17.8 / 35.6 / 46.7 |
| E | timothy grass (*Phl p 5*, *Phl p 1*, *Phl p 4*) | 22 | 50.14 (7.21) | 26.08 (3.20) | 59.1 / 40.9 | 54.5 / 27.3 / 18.2 |
| F | timothy grass (*Phl p 5*, *Phl p 1*), Bermuda grass (*Cyn d 1*) | 10 | 45.20 (4.87) | 25.94 (2.21) | 50.0 / 50.0 | 0.0 / 30.0 / 70.0 |
| G | dust mite (*Der f 2*, *Der p 2*, *Der f 1*) | 22 | 48.36 (5.14) | 26.17 (4.38) | 45.5 / 54.5 | 50.0 / 22.7 / 27.3 |
| H | Bermuda grass (*Cyn d 1*), timothy grass (*Phl p 4*, *Phl p 1*) | 40 | 51.52 (7.20) | 28.20 (5.20) | 50.0 / 50.0 | 30.0 / 37.5 / 32.5 |
| I | olive tree (*Ole e 1*), birch (*Bet v 1*), cat (*Fel d 1*) | 74 | 52.97 (5.98) | 27.20 (6.65) | 33.8 / 66.2 | 43.2 / 35.1 / 21.6 |

Demographic patterns varied across clusters. Cluster F had the youngest participants (mean age 45.2 years) and the highest proportion with higher education (70%). The oldest participants were in Cluster B (mean age 53.9 years), while body mass index was highest in Cluster H (mean 28.2). Female predominance was strongest in Cluster C (76.9%), whereas other clusters showed balanced or slightly male-skewed distributions.

To complement the clustering and latent component analysis, we also characterized cohort heterogeneity by introducing an IgE Burden Score which counts the number of detected allergen-specific IgEs per individual (Figure 1B). Among the 275 sensitized participants, 47 exhibited a single IgE response and 228 exhibited multiple responses. Based on burden, individuals were further classified into high (n = 57), medium (n = 110), and low (n = 108) scoring groups. Polysensitization profiles (”any”, “multiple”, “medium”) were dominated by birch (*Bet v 1*), timothy grass (*Phl p 1*), and Bermuda grass (*Cyn d 1*). Detailed demographic summaries for these burden-based partitions are provided in Table S1.

### Co-occurrence patterns of IgE sensitization reveal distinct clusters of pollen, house-dust mite, food, and animal dander allergens

We next characterized the cohort by pairwise co-sensitization (co-occurrence of IgEs) analysis across all 508 individuals in the cohort. We observed that most IgE pairs had low co-occurrence, with only a few exceeding 15% (Figure 2A). Sensitization clustered strongly within allergen sources, including pollen, food, house dust mites (HDM), and animal dander (see Figure 2B for a detailed breakdown), whereas cross-source co-sensitization was rare and mainly involved homologous PR-10 proteins.

**Fig. 2.**
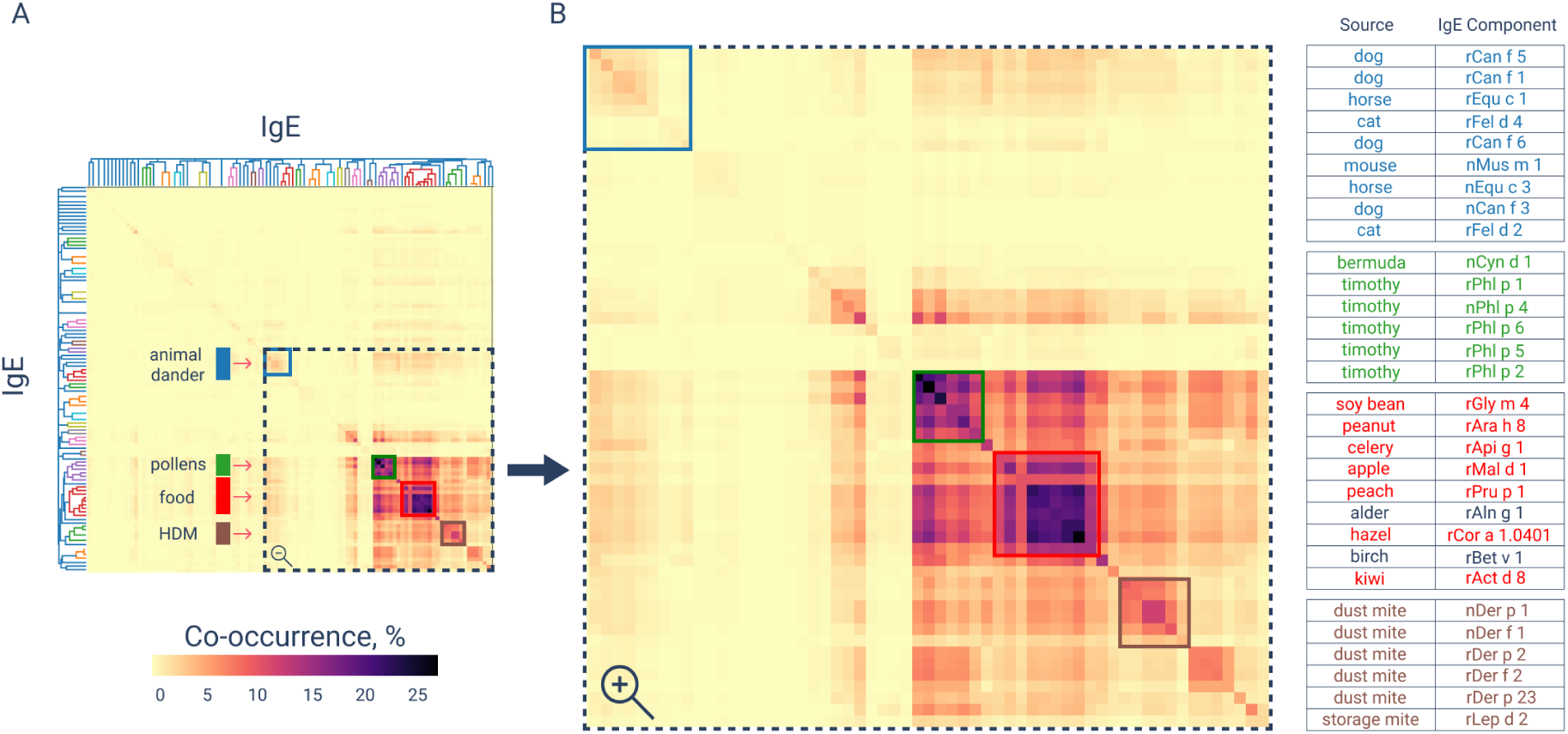
Co-occurrence analysis shows source-specific clusters of IgE sensitization(A) Heatmap of pairwise IgE co-occurrence across 508 individuals. Color intensity indicates the percentage of individuals in which two IgEs are jointly detected. Clusters of co-sensitization are annotated by allergen source: pollen (green), food (red), animal dander (blue), and HDM (brown).(B) Zoomed-in view of the co-sensitization hotspot with a table listing representative IgEs from each source-specific cluster.

A central pollen cluster (green) included timothy grass (*Phl p 1, Phl p 2, Phl p 4–6*) and Bermuda grass (*Cyn d 1*), with up to 20% co-occurrence. Additional pollen subclusters were defined by birch (*Bet v 1, v 2, v 4*), sycamore (*Pla a 1, a 3*), olive (*Ole e 1, e 7, e 9*), and alder (*Aln g 1*). A food cluster (red) comprised peanut (*Ara h 1–3, h 6, h 8, h 9*), soybean (*Gly m 4–6*), hazel (*Cor a 1.0101, a 1.0401, a 8, a 9, a 14*), peach (*Pru p 1, p 3*), apple (*Mal d 1*), and celery (*Api g 1*). Animal dander (blue) formed a separate cluster of dog (*Can f 1–6*), cat (*Fel d 1, d 2, d 4*), and horse (*Equ c 1, c 3*). HDM IgEs (brown), including *Der p 1, p 2, p 23* and *Der f 1, f 2*, clustered distinctly with strong within-source co-occurrence.

### No significant alpha diversity differences, but specific bacterial families signatures in IgE-sensitized individuals

We next examined whether the IgE-derived cohort sub-groups and partitions were associated with changes in gut microbial diversity or microbial taxa composition. For each sub-group we estimated alpha diversity using the Shannon diversity index and Faith’s Phylogenetic Diversity (Faith’s PD) from the 6,958 non-singleton ASVs present in the study cohort (see Figure S1 for the study design). We did not observe any significant differences between any of the IgE-sensitized sub-groups and non-sensitized individuals (Figure 3A). Pair-matching by sex, body mass index (BMI), and age confirmed the absence of significant differences (all p-values *>* 0.05). Contrary to prior work in food allergy cohorts that reported reduced alpha diversity in IgE-positive individuals [18], our findings indicate no microbial diversity differences in adults. Full numerical results, including sample sizes and p-values for all IgE-based partitions, are provided in Table S5.

**Fig. 3.**
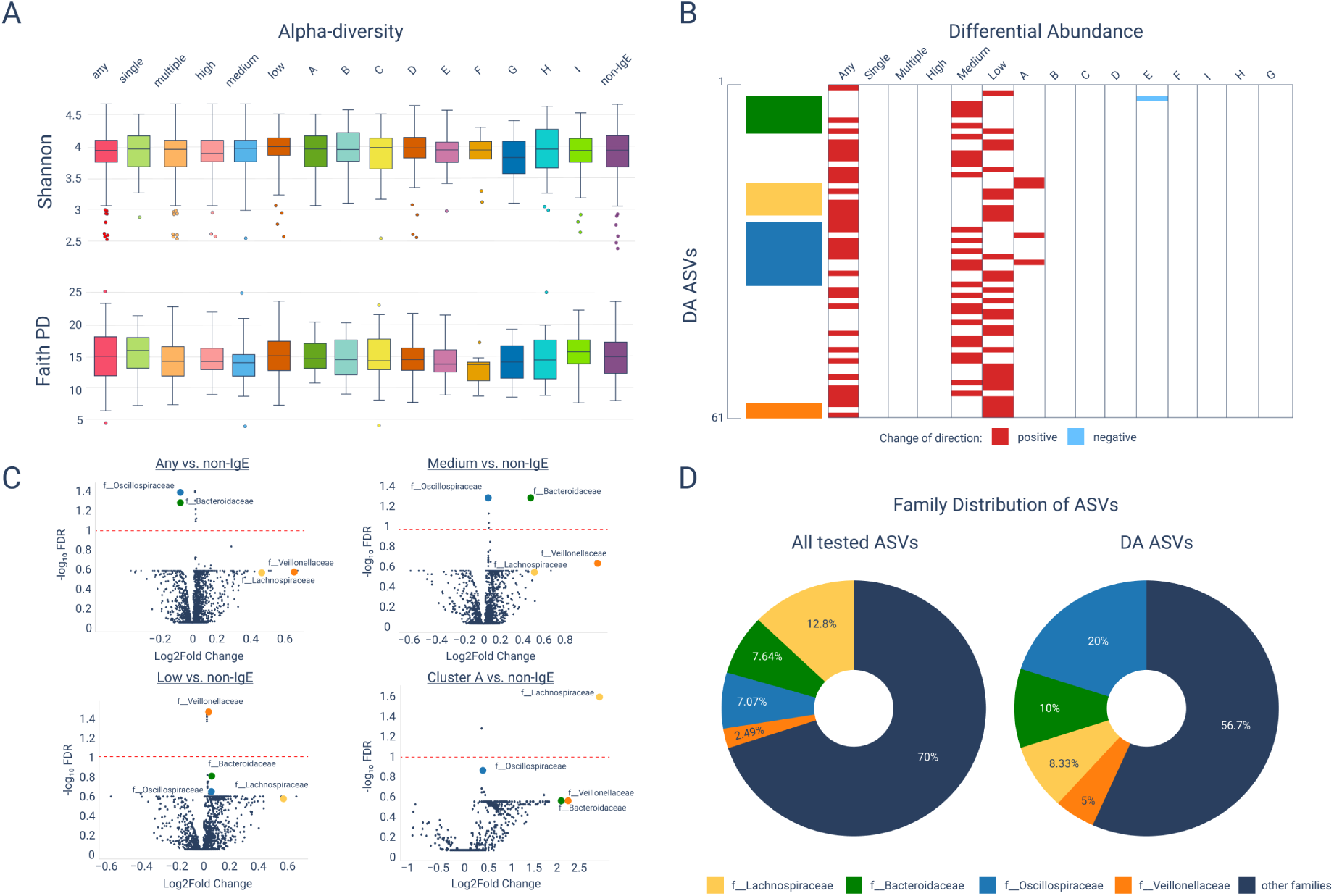
Summary of the alpha diversity and ASV-level differential abundance analysis. (A) Shannon diversity index and Faith’s Phylogenetic Diversity across IgE partitions and non-sensitized individuals. No significant differences were observed (all p-values *>* 0.05). (B) Heatmap of 61 differentially abundant ASVs across IgE partitions (red = increased, blue = decreased). (C) Volcano plots of the partitions with the strongest signals (”any”, “medium”, “low”, “Cluster A”), showing ASV-level log2 fold change and statistical significance. (D) Family-level distribution of all ASVs compared with differentially abundant ASVs. Enriched families include Bacteroidaceae (green), Lachnospiraceae (yellow), Oscillospiraceae (blue), and Veillonellaceae (orange).

Despite this lack of diversity differences, differential abundance (DA) analysis using LinDA [20] identified 61 ASVs that varied significantly across IgE partitions (Figure 3B–C). Signals were strongest in the “any”, “medium”, and “low” IgE burden groups. A few differentially abundant ASVs were also identified in Clusters A and E, respectively. When assigning family-level taxonomy to the identified ASVs, we observed that the percentage of ASVs in *Oscillospiraceae* increased from 7.1% of all ASVs to 20% of differentially abundant ASVs, ASVs in *Veillonellaceae* from 2.5% to 5%, and ASVs in *Bacteroidaceae* from 7.6% to 10%, whereas *Lachnospiraceae* decreased from 12.8% to 8.3% (Figure 3D).

Overall, these findings indicate that, while alpha diversity is not associated with IgE status, ASVs in specific microbial families, particularly *Bacteroidaceae*, *Oscillospiraceae*, and *Veillonellaceae*, are proportionally enriched, whereas ASVs in the *Lachnospiraceae* family are proportionally reduced in IgE-sensitized individuals.

### Taxonomic presence-absence analysis identifies distinct microbial taxa in IgE-sensitized and non-sensitized individuals

To complement the DA analysis, we followed recent suggestions [21, 22] and investigated the presence or absence of ASVs in IgE-sensitized or non-sensitized individuals. Specifically, we traced the proportions of ASVs in IgE-sensitized or non-sensitized individuals with (partial) taxonomic information across all taxonomic ranks (see Figure 4A and Table S2). Overall, the number of distinct ASVs was comparable between the two groups (2,280 in IgE-sensitized vs. 2,029 in non-sensitized), with 2,649 ASVs shared among both groups (Table S2). When tracing distinct ASVs through the taxonomic tree, we observed that most were concentrated in the *Clostridia* class and the orders *Coriobacteriales*, *Bacteroidales*, *Oscillospirales*, and *Rhodospirillales* (highlighted in Figure 4A). Families enriched in IgE-sensitized individuals included *Methanocorpusculaceae* and *Micrococcaceae* (highlighted in red), whereas non-sensitized individuals showed exclusive presence of families such as *Corynebacteriaceae* and *Helicobacteraceae* (highlighted in green).

**Fig. 4.**
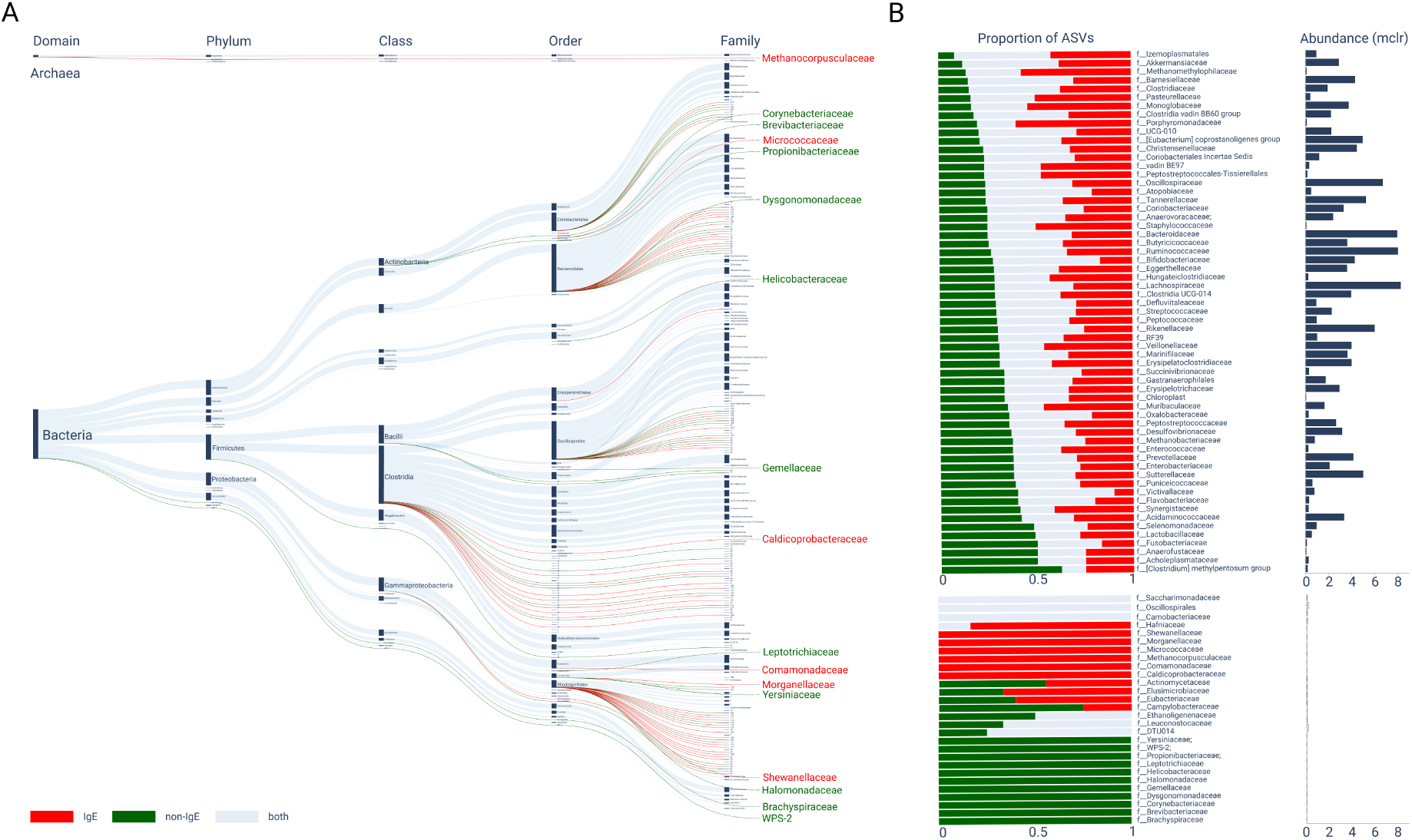
Presence–absence analysis identifies taxa distinct to IgE-sensitized or non-sensitized individuals (A) Sankey diagram showing the distribution of ASVs across taxonomic levels, with edge thickness corresponding to the number of features. (B) Proportion of ASVs within families exclusive to IgE-sensitized (red), non-sensitized (green), or shared (blue) individuals.

For the 58% of ASVs with known taxonomic family assignment, we calculated their corresponding proportional prevalence in either of the two groups (see Figure 4B, left panel). Six families comprised ASVs that only appeared in IgE-sensitized individuals (Figure 4B, full red bars), including *Shewanel-lacceae* and *Micrococcaceae*, and eleven families comprise ASVs that only appeared in non-sensitized individuals (full green bars), including *Yersiniaceae* and *Cornyebacteriaceae*. These distinct families carry, however, low relative abundance across the cohort (see Figure 4B, right panel, for corresponding modified centered log-ratio (mclr)-transformed relative abundances). While the families *Metahnomethylphilaceae*, *Monoglobaceae*, *Porphyromonadaceae*, *Staphylococcaceae*, and *Hafniaceae* comprised more than 50% ASVs in IgE-sensitized individuals, the families *Clostridium methylpentosum group*, *Actinomycetaceae*, and *Campylobacteraceae* comprised more than 50% distinct ASVs in non-sensitized individuals. None of these families are highly abundant in the cohort, thus making it difficult to be detected by compositional DA analysis. Conversely, none of the families that were enriched for DA ASVs showed a particularly skewed prevalence in either of the groups. Nevertheless, our taxonomic presence-absence analysis provides a more fine-grained view of family prevalence across IgE-sensitized and non-sensitized individuals that may help generate future functional hypotheses.

### Limited predictability of IgE profiles from microbial taxa abundances

We next investigated whether microbial relative abundances could be used to distinguish the nine clustered sub-groups and six IgE burden-based partitions from non-sensitized individuals. Given the limited sample size of most sub-groups and partitions, we performed classification analysis using compositionally-aware penalized log-contrast classification [23, 24]. This approach allows to identify a small set of log-ratios of microbial taxa that best predict class membership. To select the right level of sparsity and guard against overfitting, we used three different model selection strategies for the penalty parameter *λ*: (i) a theoretically-derived penalty parameter (*λ*_fixed_) [25, 26], (ii) 5-fold cross-validation (*λ*_CV_), and (iii) stability selection (*λ*_SS_) [26]. The latter method proves to be particularly beneficial for microbiome data [27, 28] and enables automatic identification of a small stable set of predictive taxa. We performed classification across all seven taxonomic ranks, resulting in 315 models (15 classes × 7 ranks × 3 model selection schemes).

Balanced out-of-sample mis-classification rates (MCRs) varied between 0.25 and 0.75. Models built at the family or genus level generally achieved lower MCRs compared to those at higher (phylum/class) or lower (species/ASV) ranks (see Supplementary Tables S3–S4). Family- and genus-rank models also tended to be more parsimonious, selecting fewer taxa while retaining comparable predictive accuracy. While the overall performance suggests that microbial abundances alone are not predictive for most IgE profiles, stability selection revealed on each taxonomic level a small set of taxa that were robustly included across all classification models. Figure 5A summarizes stability results on family level. Here, we observed that several families, including *Veillonellaceae*, *Ruminococcaceae*, *Rikenellaceae*, *Prevotellaceae*, *Oscillospiraceae*, *Lachnospiraceae*, *Christensenellaceae*, and *Bacteroidaceae* were selected with high stability (i.e., had high selection probability of being included as a predictor) across most partitions. Notably, four of these families (colored in Figure 5A) also appeared in the differential abundance analysis, underscoring their potential association with IgE status. In contrast, families such as *Akkermansiaceae* and *Eggerthellaceae* were selected less frequently and appeared to be partition-specific.

**Fig. 5.**
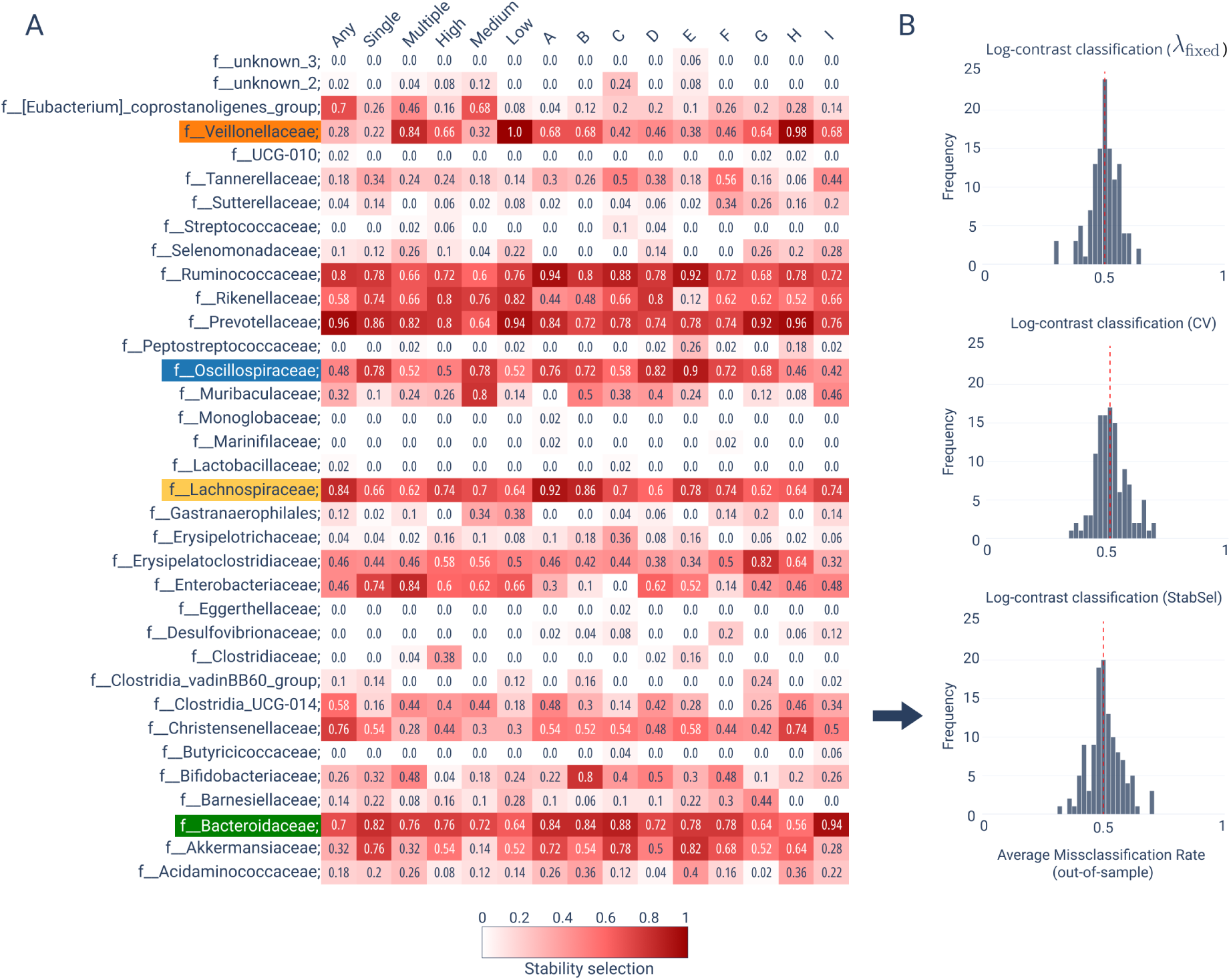
Summary of log-contrast classification of IgE sensitization patterns from microbial relative abundances. (A) Selection probabilities of each microbial family (rows) for inclusion in log-contrast classification models using stability selection for model selection across all 15 IgE partitions and clusters (columns). Darker red color indicates higher stability selection probability. Names of the four families that comprised a higher proportion of DA ASVs are highlighted in color. (B) Histograms of out-of-sample misclassification rates (MCRs) for all 105 classification models (seven taxonomic ranks across 15 IgE partitions and clusters) for three different model selection strategies. The top histogram shows MCRs using a theoretically derived penalization parameter *λ*_fixed_, the middle histogram 5-fold CV with *λ*_CV_, and the lower histogram stability selection with *λ*_7tab7el_. The dashed red line indicates a median MCR of 0.5.

The choice of the model selection strategy for the penalty parameter *λ* in the sparse log-contrast models had a negligible effect on overall classification performance, as shown in Figure 5B. Across the 315 models, each model selection scheme produced similar out-of-sample MCR distributions with a median rate of 0.5 (see Table S3 and S4 for individual results).

Taken together, our comprehensive classification analysis shows that microbial relative abundances cannot predict IgE status, independent of the taxonomic aggregation level. Stability analysis, however, identified small sets of taxa that are reproducibly selected as predictors for classification, with family-and genus-level features offering the best balance between accuracy and interpretability (see Table S3 and S4 for comparison).

### Network analysis reveals IgE-specific alterations in microbial family associations

We next sought to identify altered taxon-taxon association patterns between (sub-groups of) IgE-sensitized and non-sensitized individuals. For ease of exposition, we agglomerated the ASV data to family rank and focused on associations among the most abundant microbial families in each sub-group. We estimated sparse partial correlation networks with SPIEC-EASI [29] and used NetCoMi [30] for network comparison and visualization. Figure 6 shows the derived networks among all families with at least one association across the respective comparisons (left panel control, middle panel sub-group, right panel difference network).

**Fig. 6.**
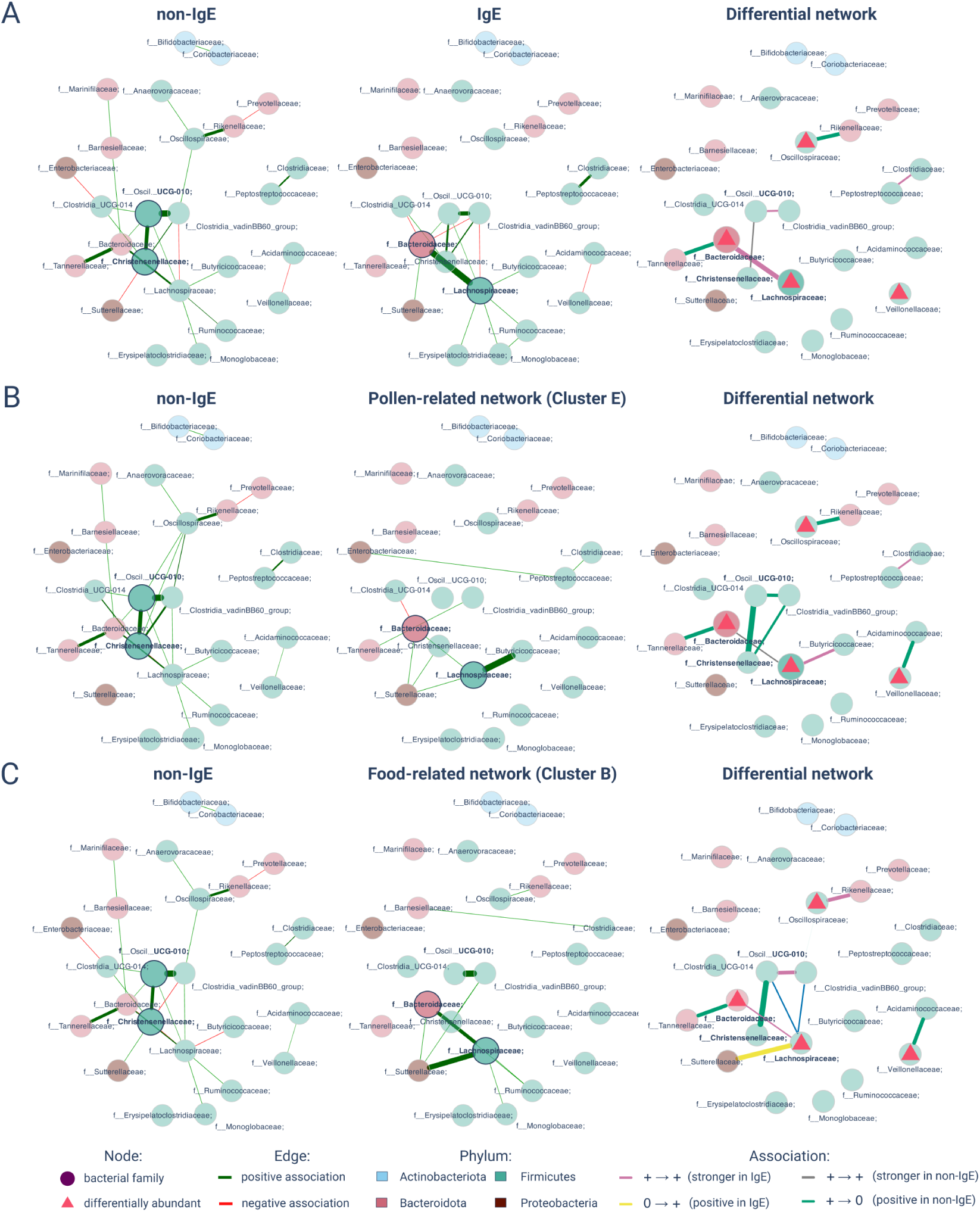
Microbial associations analysis (A–C) Left: non-IgE networks; middle: IgE-partitions networks (A: any-IgE; B: pollen-related Cluster E; C: food-related Cluster B); right: differential networks showing non-IgE to IgE changes. Edge thickness reflects association strength (green = positive, red = negative); node color indicates phylum; triangles mark differentially abundant families. Differential edge colors: pink (positive to positive, stronger in IgE), grey (positive to positive, stronger in non-IgE). yellow (absent to positive, only IgE), green (positive to absent, only non-IgE).

To get a global picture, we first estimated and compared association networks between 42 microbial families derived from *n* = 233 non-sensitized individuals (non-IgE) and 41 microbial families from the *n* = 275 individuals with at least one IgE marker present (IgE; Figure 6A). The non-IgE networks shows a large connected component comprising 19 families with *Bacteroidaceae*, *Christensenellaceae*, *Lachnospiraceae*, and *Oscillospirales UCG-010* forming the core of the network component. All but five associations are estimated to be positive. In the IgE network, the largest connected component comprises eleven families. Stronger associations were estimated between *Bacteroidaceae* and *Lachnospiraceae* and *Peptostrepococcaceae* and *Clostridiaceae*. Weaker association were estimated between *Oscillospirales UCG-010* and *Christensenellaceae*. No positive associations were estimated between *Bacteroidaceae* and *Tannerellaceae*, and *Oscilliospiraceae* and *Rikenellaceae*, respectively. A summary of the differential network, highlighting key changes in associations are summarized in Figure 6A, right panel. Here, we also marked the families that comprised differentially abundant ASVs with red triangles: *Veillonellaceae*, *Bacteroidaceae*, *Lachnospiraceae*, and *Oscilliospiraceae*. For the latter three, these differentially abundant ASVs may contribute to the observed differences in global associations.

We next focused on network analysis for two important sub-groups of IgE-sensitized individuals: (i) the pollen-related subgroup Cluster E (n=22 individuals, p=38 microbial families) that is characterized by high loadings only in latent allergy component 2 (LAC2), and (ii) food-related subgroup Cluster B (n=24 individuals, p=43 microbial families) that show high loadings in latent allergy component 1. We compared the association networks derived from these specific subgroups with networks derived from random subsets of non-IgE (control) individuals of corresponding size. Figure 6B and Figure 6C summarize the resulting association networks and their key differences. We observed few differences between the respective (sub-sampled) non-IgE networks, both exhibiting large nearly identical connected components. Differences to the global non-IgE network simply reflects the reduced sample size at construction. For the sub-group specific networks, however, we observed marked differences in association patterns. For the poller-related network (Figure 6B, middle panel), we observed a stronger positive association between *Lachnospiraceae* and *Butyricicoccaceae* families. All other associations were reduced in strength or reversed sign. For the food-related network (Figure 6C, middle panel), we observed considerable rewiring around the *Lachnospiraceae* and *Bacteroidaceae* families. *Lachnospiraceae* showed a stronger positive association with *Bacteroidaceae* and formed a new association with *Sutterellaceae*. *Bacteroidaceae* on the other hand lost most of its associations and formed a new one with *Sutterellaceae*. Additionally, the strong association between *Oscillospirales UCG-010* and *Christensenellaceae* vanished in the food-related network.

Taken together, our analysis revealed altered network signatures both on the global level (IgE vs. non-IgE) and on sub-group level, as illustrated with the pollen- and food-related networks. Integration of these network results with our prior LinDA-based DA analysis revealed both concordant and complementary patterns. Families enriched with differentially abundant ASVs, such as *Oscillospiraceae*, *Veillonellaceae*, *Lachnospiraceae*, and *Bacteroidaceae*, participated in network rewiring but could not fully explain the estimated differences.

### Taxon set enrichment links vitamin- and metabolite-producing taxa to IgE sensitization

To complement our prior taxonomy-focused analysis, we next sought to understand whether certain sets of species or genera with specific metabolic functions or predicted metabolic potential are altered in IgE-sensitized and non-sensitized individuals. To perform this analysis, we used the recently introduced Taxon set enrichment analysis (TaxSEA) framework [31] which integrates taxon sets with known metabolic capabilities from several public microbiota databases. Analogous to popular Gene Set Enrichment Analysis [32] in genomics, TaxSEA groups sets of species or genera (with known taxonomic assignment) according to function and computes group enrichment scores from the corresponding log-fold changes between different conditions.

Here, we performed TaxSEA on the *p* = 229 species with known taxonomic assignment to identify potential enrichment among IgE-sensitized (*n* = 275) and non-sensitized individuals (*n* = 233). We identified 21 metabolite-related sets associated with IgE sensitization, including groups linked to short-chain fatty acids (SCFAs), bile acids, amino acid derivatives, and vitamins. Among these, folic acid (99 taxa) and vitamin A (34 taxa) sets were most significantly enriched (Figure 7A; Figure S2A; Table S6-S7).

**Fig. 7.**
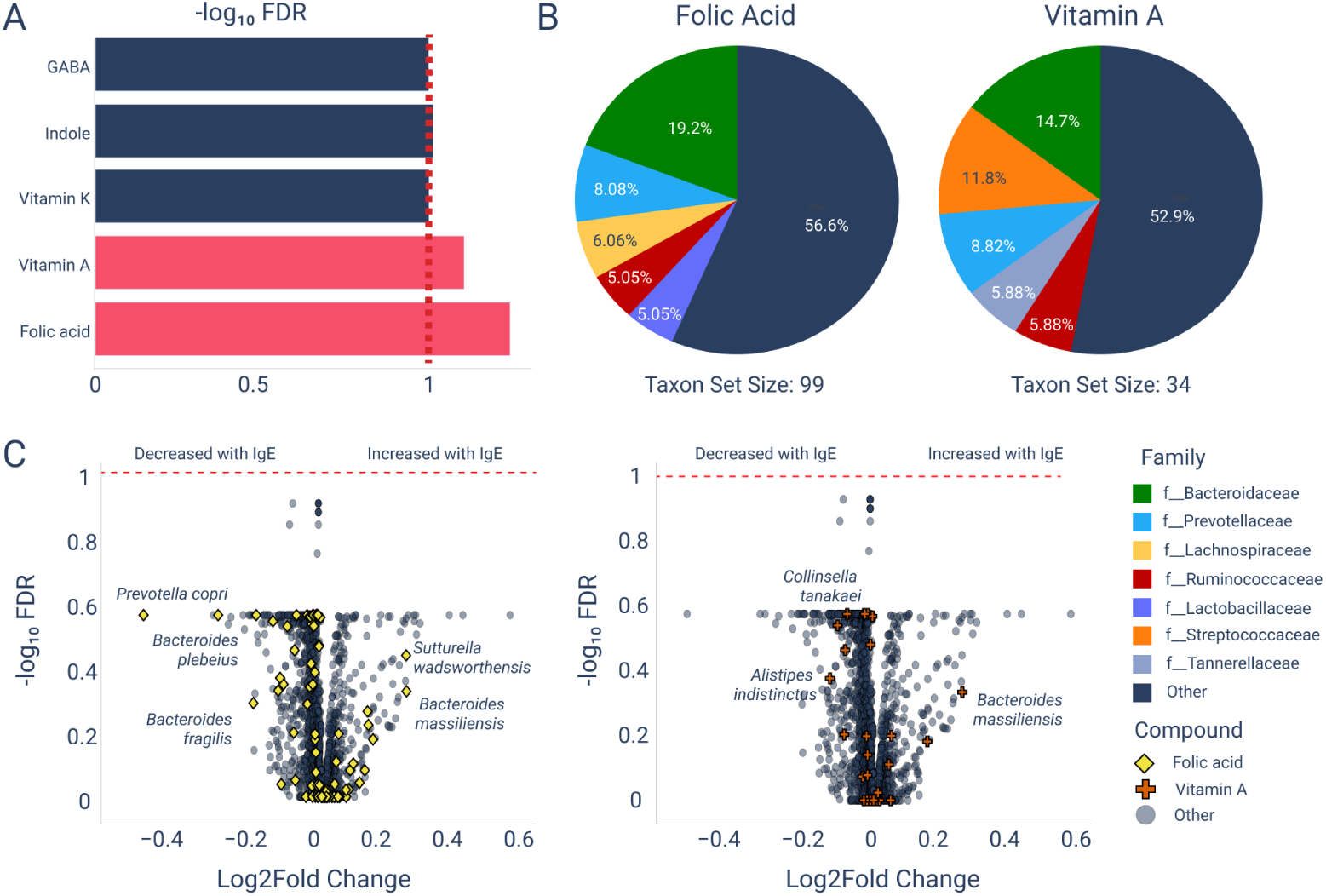
Taxon set enrichment analysis reveals vitamin- and metabolite-producing taxa associated with IgE sensitization (A) Significantly enriched taxon sets (FDR *≤* 0.1). (B) Family-level composition of folic acid and vitamin A taxon sets (top five families shown). (C) Volcano plots of species-level abundance changes for folic acid and vitamin A producer sets.

Family-level composition of these sets was dominated by *Bacteroidaceae*, with contributions from *Prevotellaceae*, *Lachnospiraceae*, *Ruminococcaceae*, *Lactobacillaceae*, *Streptococcaceae*, and *Tannerellaceae* (Figure 7B). This indicates that a small number of families account for the majority of vitamin-producing taxa. Consistent with prior reports of reduced SCFA producers in allergy [18], we observed decreased abundance of *Prevotella copri*, *Eubacterium ramulus*, and *Bacteroides massiliensis* in IgE-sensitized individuals.

At the species level, several folic acid producers, including *Prevotella copri*, *Bacteroides plebeius*, and *Bacteroides fragilis*, were depleted, whereas *Sutturella wadsworthensis* and *Bacteroides massiliensis* increased (Figure 7C). Within the vitamin A producer set, *Alistipes indistinctus* and *Collinsella tanakaei* decreased modestly, while *Bacteroides massiliensis* showed a strong positive shift. Notably, the opposite responses of *Prevotella copri* (depletion) and *Bacteroides massiliensis* (enrichment) recurred across multiple metabolite sets, suggesting they are consistent drivers of functional differences between IgE-positive and IgE-negative individuals.

Together, these results highlight folic acid- and vitamin A-producing taxa, and, in particular, the opposing roles of *Prevotella copri* and *Bacteroides massiliensis*, as potential microbial signatures linked to IgE sensitization.

## 3 DISCUSSION

In this contribution, we have presented a multimodal large-scale integrative analysis of deep allergen-specific IgE profiles and gut microbiome amplicon sequencing data from 508 adults in the population-based KORA FF4 cohort. Our primary objective was to identify potential relationships between IgE sensitization patterns and gut microbial features in a quantitative manner. Despite widespread consensus that the gut microbiome plays a contributing role in allergic disease development, manifestation, and attenuation [12], few studies have been available that jointly analyze IgE and gut microbiome patterns on cohort scale [18].

We have tackled this challenge by first identifying statistically meaningful and interpretable sub-groups (or strata) of individuals from the available deep allergen-specific IgE profiles. Using techniques from hierarchical clustering, dimensionality reduction [19], and co-occurrence analysis, we have identified eight sub-groups (clusters) of IgE-sensitized individuals with broadly similar patterns as well as one group with sparse unspecific IgE patterns (Figure 5; Cluster I). Notably, we could also show through careful dimensionality reduction analysis that three latent allergy components (LACs) suffice to explain the observed IgE sensitization patterns in the adult cohort. IgE co-occurrence analysis confirmed that these components largely reflect allergen sources, including pollen, food, house dust mites, and animal dander, an observation consistent with prior large-scale analyses [7, 9]. Such reproducible patterns highlight the influence of environmental exposures and molecular cross-reactivity on sensitization profiles, particularly within the PR-10 protein family. These findings support the value of stratifying individuals by allergen-specific IgE, and provide a principled basis for examining IgE sensitization relationships with other data modalities within and across cohort sub-groups [2].

However, relating the identified sub-groups to the available large-scale microbial abundance patterns proved to be challenging. For instance, we have not been able to detect significant alpha diversity changes between sensitized and non-sensitized individuals irrespective of the considered sub-groups, and independent of overall IgE burden. This indicates that previously observed patterns of significant microbial diversity changes [18] cannot be reproduced in the present adult cohort. This also suggests that alpha diversity shifts may be context-dependent, e.g., occurring in individuals with specific food allergies, and are not generalizable to broader adult populations. Notwithstanding these population-level results, we have been able to identify sets of amplicon sequence variants (ASVs) across multiple cohort partitions that showed statistically significant differences. When aggregated on family-level, these ASVs were overrepresented in *Bacteroidaceae*, *Oscillospiraceae*, and *Veillonellaceae* and underrepresented in *Lachnospiraceae*. This may potentially hint at a shift in the balance between mucin-degrading and barrier-protective taxa implicated in allergic sensitization [15, 16]. For completeness, we have also performed a strain-level presence-absence analysis to identify potentially distinct patterns in IgE-sensitized and non-sensitized individuals [22]. While we have been able to identify unique ASVs in the respective groups, their overall relative abundances were low, suggesting that deeper amplicon profiling or larger cohorts are necessary to discover robust strain-level biomarkers. Similarly, microbial relative abundances have turned out to provide only limited predictability of IgE patterns. Despite using dedicated classification schemes for high-dimensional relative abundance data [25–27], we have not been able to reliably predict IgE sub-group membership from microbiome data alone, independent of taxonomic rank aggregation and model selection strategy. This adds further evidence that microbial abundance measurements from amplicon sequencing may be insufficient to capture intricate patterns of allergy sensitization.

Our microbial network analysis, on the other hand, identified notable family-family association differences in IgE-sensitized individuals with respect to *Lachnospiraceae*, *Bacteroidaceae*, and *Chris-tensenellaceae* forming the central core of the community structure. When focusing on specific sub-groups of IgE-sensitized individuals, including the pollen-related and the food-related subgroups, we discovered condition-specific network differences, most prominently involving associations with *Lachnospiraceae*. Whilethe biological interpretation of these network-level changes remains preliminary, we have shown that statistical association networks can reveal differences that would have remained undetected in mere diversity or differential abundance analysis. Finally, we have introduced a framework fo functional enrich-ment analysis in the context of IgE sensitization. Using recent tools from taxon set enrichment analysis [31] we have been able to link IgE status to taxa producing folic acid and vitamin A. Several species within these sets, including *Prevotella copri* and *Bacteroides massiliensis*, showed opposite associations that recurred across multiple metabolite groups. In the absence of metabolomics data for the present cohort, and given that our analysis relied on 16S rRNA-based predictions rather than direct metabolite measurements, these findings should be interpreted as hypothesis-generating. Nevertheless, they are consistent with prior reports of significantly higher SCFA levels in non-allergic controls [18], suggesting a relationship between IgE status and the metabolic profile of the gut microbiome.

Taken together, by integrating deep allergen-specific IgE profiles, microbiome data, and epidemiological data within the well-characterized KORA FF4 cohort, our study represents one of the most comprehensive cross-sectional analyses of the gut–immune axis in IgE-mediated allergy in adults to date. For these data, we have introduced a dedicated statistical analysis framework, resulting in several testable hypotheses in the form of microbial taxa, families, and metabolite producers that may play a role in IgE sensitization in adults.

While this contributes to ongoing efforts to understand the relationship between the gut microbiome and IgE-mediated allergy, several limitations must be acknowledged. Firstly, our results should be interpreted cautiously since the cross-sectional design naturally excludes any causal statements. Whether specific microbial configurations actively drive IgE sensitization, whether immunological predisposition shapes the gut microbiome, or whether both reflect a shared underlying allergic predisposition remains unresolved. Secondly, for the current study, neither full metagenomics sequencing data nor metabolomics data have been available. Given the observed limited predictability of amplicon data alone, a full integration of deep IgE profiling with metagenomics and metabolomics data may be warranted. Thirdly and most critically, dietary data were not available for the present cohort, which represents an important confound. For instance, under conditions of low dietary fiber, a characteristic of Western diets, mucin-degrading bacteria shift from consuming dietary substrates to foraging on host mucus, thinning the intestinal barrier and potentially facilitating direct contact between luminal allergens and immune cells [15, 33]. While we can only speculate that this diet–microbiota–mucus axis may partly underlie the microbial signatures we observe, e.g., the enrichment of *Bacteroidaceae*, the relative depletion of *Lachnospiraceae*, and altered association networks, future studies should include dietary covariates to further disentangle these intricate relationships. Finally, as the field moves forward, we posit that longitudinal sampling, mechanistic experiments in gnotobiotic models, and causal inference approaches such as Mendelian randomization, are necessary to decipher the complex relationship between IgE-mediated allergy and the microbiome.

## RESOURCE AVAILABILITY

### Lead contact

Requests for further information and resources should be directed to and will be fulfilled by the lead contact, Oleg Vlasovets.

### Materials availability

This study did not generate new unique reagents.

### Data and code availability

- 16S rRNA gene sequencing and allergen-specific IgE data from the KORA FF4 cohort cannot be deposited in public repositories due to participant consent restrictions. Data can be requested for research projects via the KORA.PASST use and access hub: https://helmholtz-muenchen.managed-otrs.com/external.
- All original code for analysis and visualization is available on GitHub at: https://github.com/bio-datascience/ige-gut-microbiome-analysis. A Docker image containing the computational environment used for this study is publicly available on DockerHub: https://hub.docker.com/repository/docker/ovlasovets/rpy2/general.
- Any additional information required to reanalyze the data reported in this paper is available from the lead contact upon request.

## ACKNOWLEDGMENTS

We thank all KORA study participants for their long-term commitment, the study staff for data collection and research data management, and the members of the KORA Study Group (https://www.helmholtz-munich.de/en/epi/cohort/kora) who are responsible for the study design and conduct. We thank Ayse Sener and Aline Metz for their excellent technical assistance with the ISAC112 assays.

## AUTHOR CONTRIBUTIONS

Conceptualization, O.V., C.L.M., M.S., and A.P.; Methodology, O.V. and C.L.M.; Software, O.V.; Formal Analysis, O.V. and C.L.M.; Investigation, O.V. and C.L.M.; Resources, C.L.M. and A.P..; Data Curation, L.M., S.G., H.G., C.H. and KORA Study Group; Writing – O.V.,C.L.M.; Review & Editing, O.V., M.S., L.M, C.L.M. and A.P..; Visualization, O.V.; Supervision, C.L.M., M.S. and A.P.; Funding Acquisition, C.L.M. and A.P..

## DECLARATION OF GENERATIVE AI AND AI-ASSISTED TECHNOLOGIES

During the preparation of this work, O.V. used **Grammarly (Superhuman Platform Inc.)** and **ChatGPT (OpenAI)** for **language editing and formatting assistance only**. The authors reviewed and edited all content and take full responsibility for the manuscript.

## SUPPLEMENTAL INFORMATION INDEX

**Document S1.** Figures S1–S2 and Tables S1–S8, plus Supplemental References

**Fig. S1.**
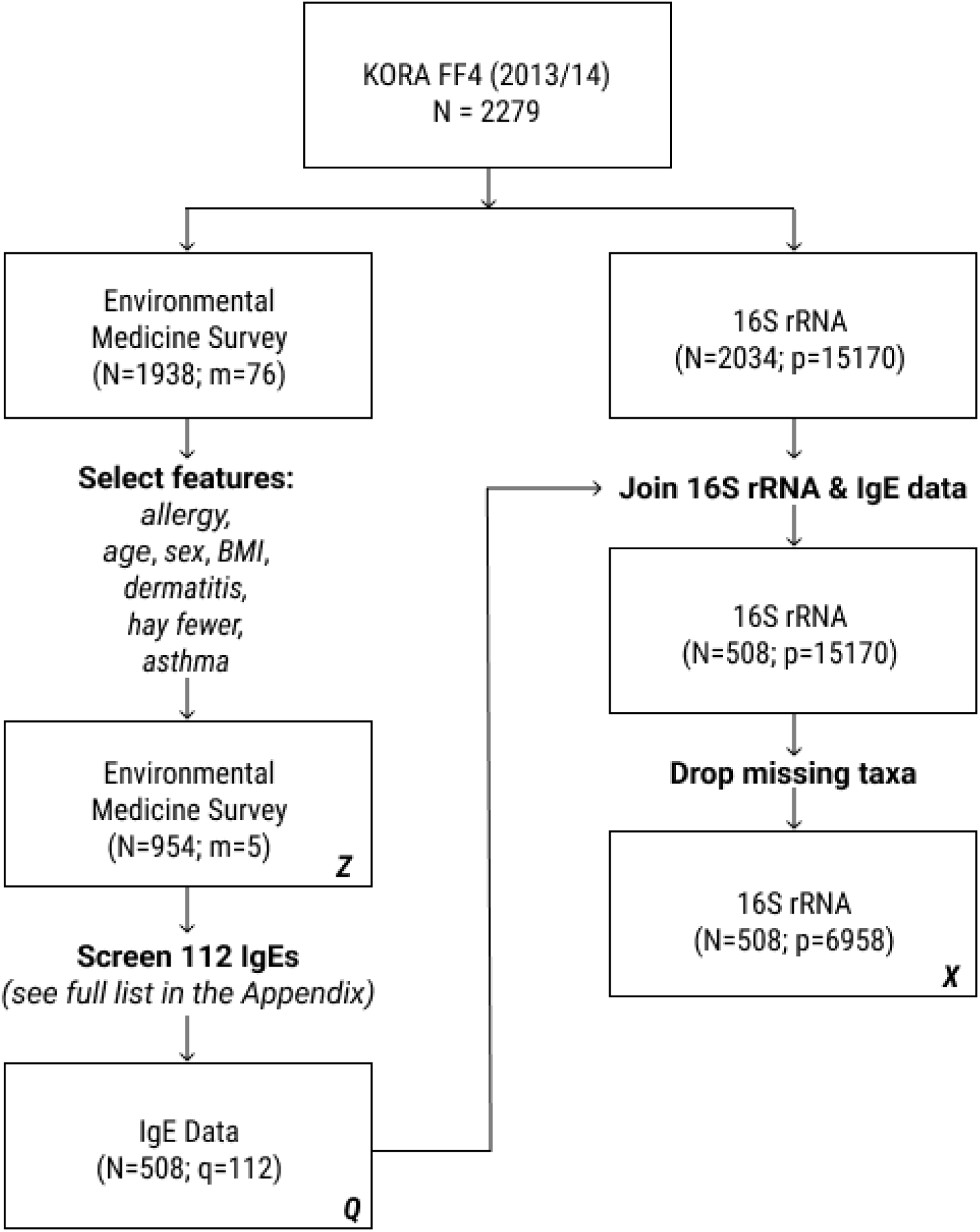
Flowchart of study design and data integration. Participant flow in the KORA FF4 study, from the initial cohort (N = 2,279) to the final analytic datasets. After survey filtering, 910 individuals (Z) were retained; IgE screening yielded 508 with complete profiles for 112 markers (Q). 16S rRNA sequencing data from 2,034 individuals were aligned with IgE data, resulting in 508 participants with 6,958 microbial features (X).

**Fig. S2.**
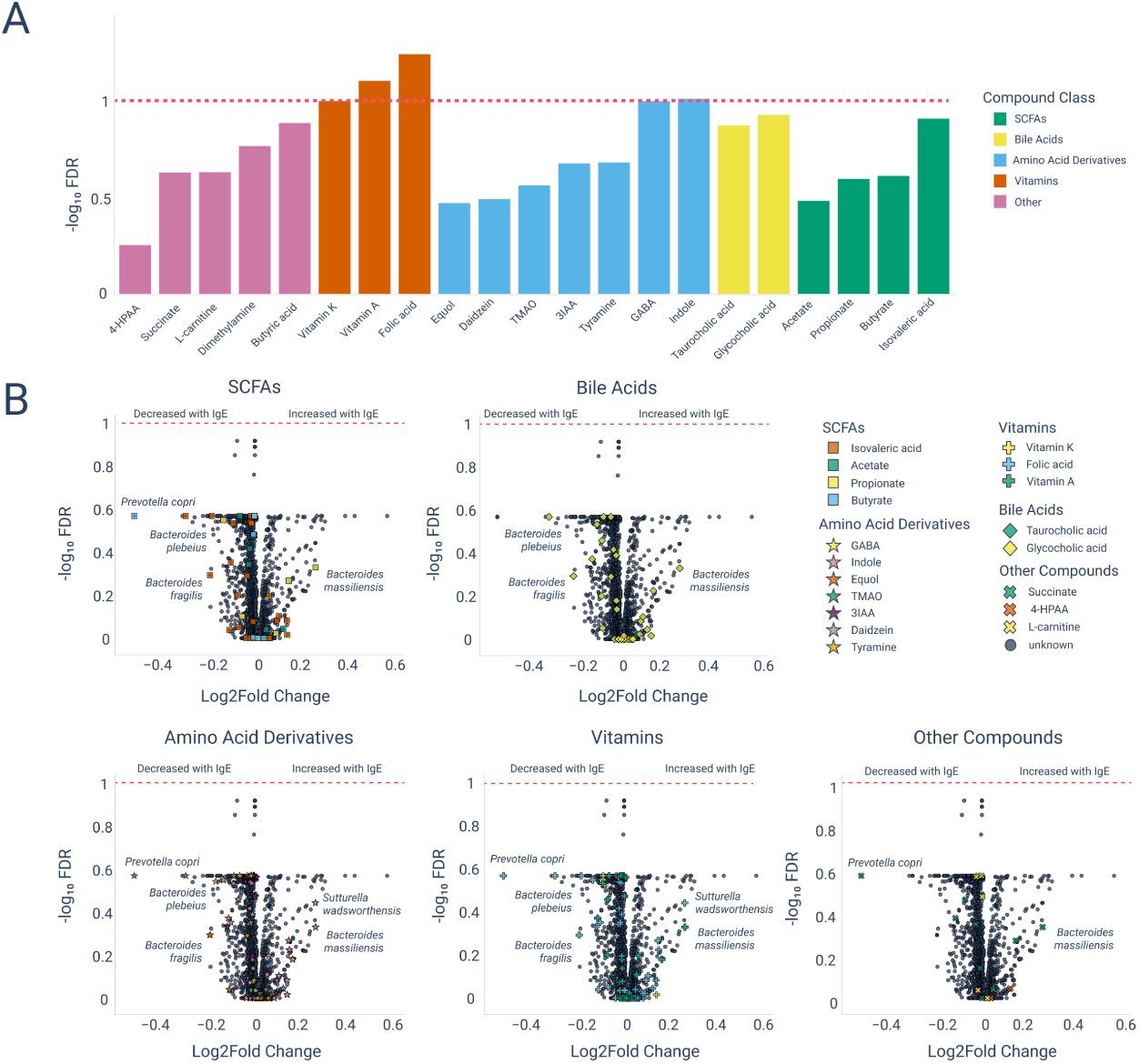
Taxon Set Enrichment Analysis (TaxSEA) of microbial metabolite-related functions.(A) Enriched taxon sets across compound classes identified by TaxSEA. Bars are colored by compound class, with the horizontal dashed line indicating the significance threshold (FDR *≤* 0.1). (B) Volcano plots of species-level differential abundance for each compound class. Points represent individual species, with select taxa labeled. Notable species such as Prevotella copri, Bacteroides plebeius, Bacteroides fragilis, and Bacteroides massiliensis are highlighted for their consistent associations across multiple compound classes.

**Table S1.**
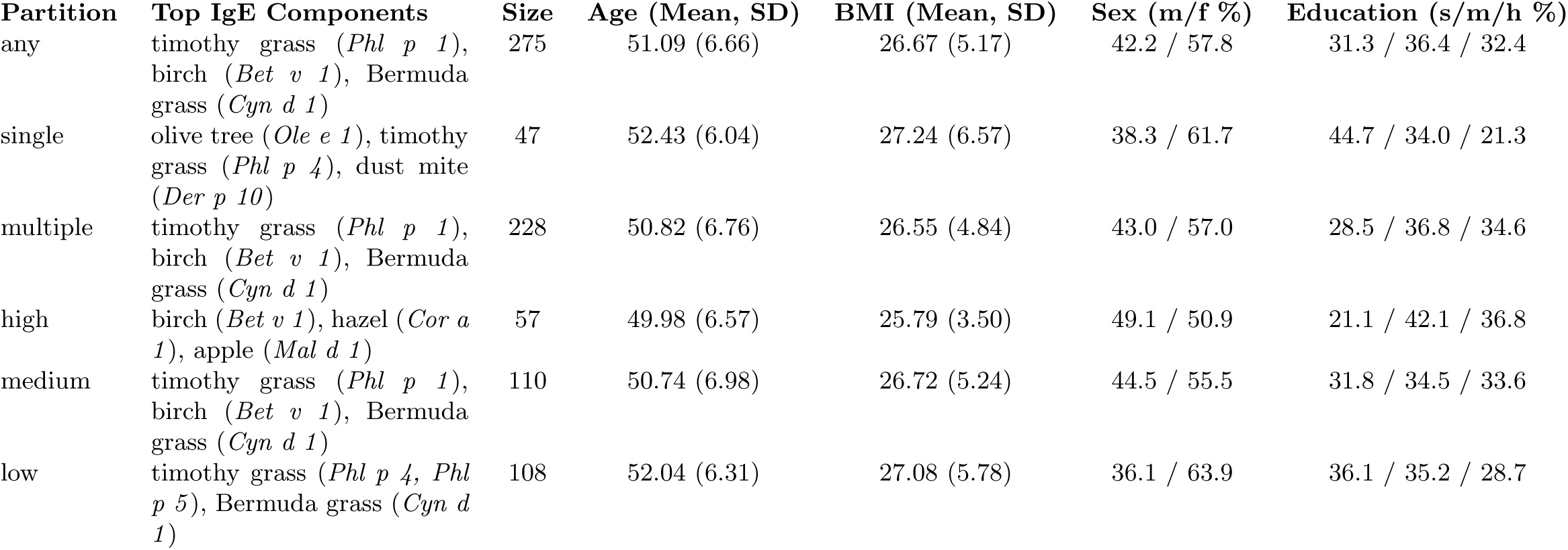
Summary statistics of IgE data partitions including age, BMI, cluster size, and the most prevalent allergen-specific IgEs. Sex distribution is shown as percentage male/female, and education is reported as percentage with secondary (s), medium (m), and higher (h) education.

**Table S2.** Distinct microbial taxa across IgE-sensitized and non-sensitized individuals at different taxonomic ranks. Counts include exclusive and shared taxa, total numbers, and proportion lacking taxonomic annotation.

| Taxonomic rank | IgE | non-IgE | IgE ∪ non-IgE | Total number of taxa | Unknown taxonomy (%) |
| --- | --- | --- | --- | --- | --- |
| Phylum | 1 | 2 | 14 | 17 | 0% |
| Class | 1 | 3 | 22 | 26 | 8% |
| Order | 11 | 14 | 62 | 87 | 40% |
| Family | 48 | 49 | 115 | 212 | 58% |
| Genus | 364 | 345 | 636 | 1,345 | 81% |
| Species | 2,038 | 1,822 | 2,442 | 6,302 | 96% |
| ASV | 2,280 | 2,029 | 2,649 | 6,958 | — |

**Table S3.** Out-of-sample misclassification rate (MCR) values for IgE frequency and score groups across taxonomic levels.

| Level | Any IgE (N=275) | Single (N=47) | Multiple (N=228) | High Score (N=57) | Medium Score (N=110) | Low Score (N=108) |
| --- | --- | --- | --- | --- | --- | --- |
| **Theoretical $\lambda_{\text{fixed}}$** |  |  |  |  |  |  |
| Phylum | 0.49 (4) | 0.61 (3) | 0.6 (5) | 0.5 (2) | 0.45 (6) | 0.5 (7) |
| Class | 0.48 (5) | 0.56 (2) | 0.59 (5) | 0.5 (4) | 0.48 (8) | 0.57 (9) |
| Order | 0.51 (6) | 0.56 (3) | 0.58 (12) | 0.5 (4) | 0.5 (7) | 0.57 (12) |
| Family | 0.44 (3) | 0.67 (6) | 0.47 (6) | 0.55 (4) | **0.38** (11) | 0.6 (6) |
| Genus | 0.49 (4) | 0.61 (4) | 0.45 (10) | 0.55 (2) | 0.4 (18) | 0.6 (7) |
| Species | 0.52 (19) | 0.5 (24) | 0.41 (24) | 0.41 (14) | 0.43 (27) | 0.55 (11) |
| ASVs | 0.59 (55) | 0.56 (19) | 0.53 (56) | 0.41 (12) | 0.5 (27) | 0.52 (18) |
| **Cross-validation** |  |  |  |  |  |  |
| Phylum | 0.5 (2) | 0.61 (2) | 0.58 (7) | 0.5 (2) | 0.64 (2) | 0.5 (7) |
| Class | 0.49 (2) | 0.56 (2) | 0.5 (5) | 0.5 (10) | 0.5 (0) | 0.57 (6) |
| Order | 0.5 (2) | 0.56 (2) | 0.56 (18) | 0.5 (12) | 0.55 (2) | 0.57 (16) |
| Family | 0.46 (3) | 0.56 (2) | 0.46 (7) | 0.55 (4) | 0.5 (0) | 0.55 (7) |
| Genus | 0.49 (3) | 0.56 (6) | 0.44 (11) | 0.45 (9) | 0.5 (0) | 0.62 (9) |
| Species | 0.5 (13) | 0.72 (6) | 0.45 (36) | **0.23** (25) | 0.45 (5) | 0.45 (2) |
| ASVs | 0.52 (25) | 0.61 (2) | 0.49 (49) | 0.36 (16) | 0.45 (173) | 0.5 (3) |
| **Stability Selection** |  |  |  |  |  |  |
| Phylum | 0.51 (5) | 0.61 (2) | 0.55 (5) | 0.55 (3) | 0.5 (6) | 0.5 (6) |
| Class | 0.43 (5) | 0.56 (2) | 0.62 (6) | 0.5 (2) | 0.52 (4) | 0.52 (7) |
| Order | 0.57 (5) | 0.56 (3) | 0.54 (7) | 0.5 (3) | 0.48 (4) | 0.6 (7) |
| Family | 0.47 (5) | 0.67 (5) | 0.49 (5) | 0.55 (4) | 0.4 (5) | 0.57 (4) |
| Genus | 0.49 (2) | 0.5 (2) | 0.45 (3) | 0.55 (3) | 0.5 (2) | 0.5 (3) |
| Species | 0.61 (3) | 0.5 (0) | 0.45 (1) | 0.45 (5) | **0.31** (2) | 0.5 (3) |
| ASVs | 0.5 (0) | 0.5 (0) | 0.5 (1) | 0.5 (4) | 0.57 (1) | 0.55 (2) |

**Table S4.**
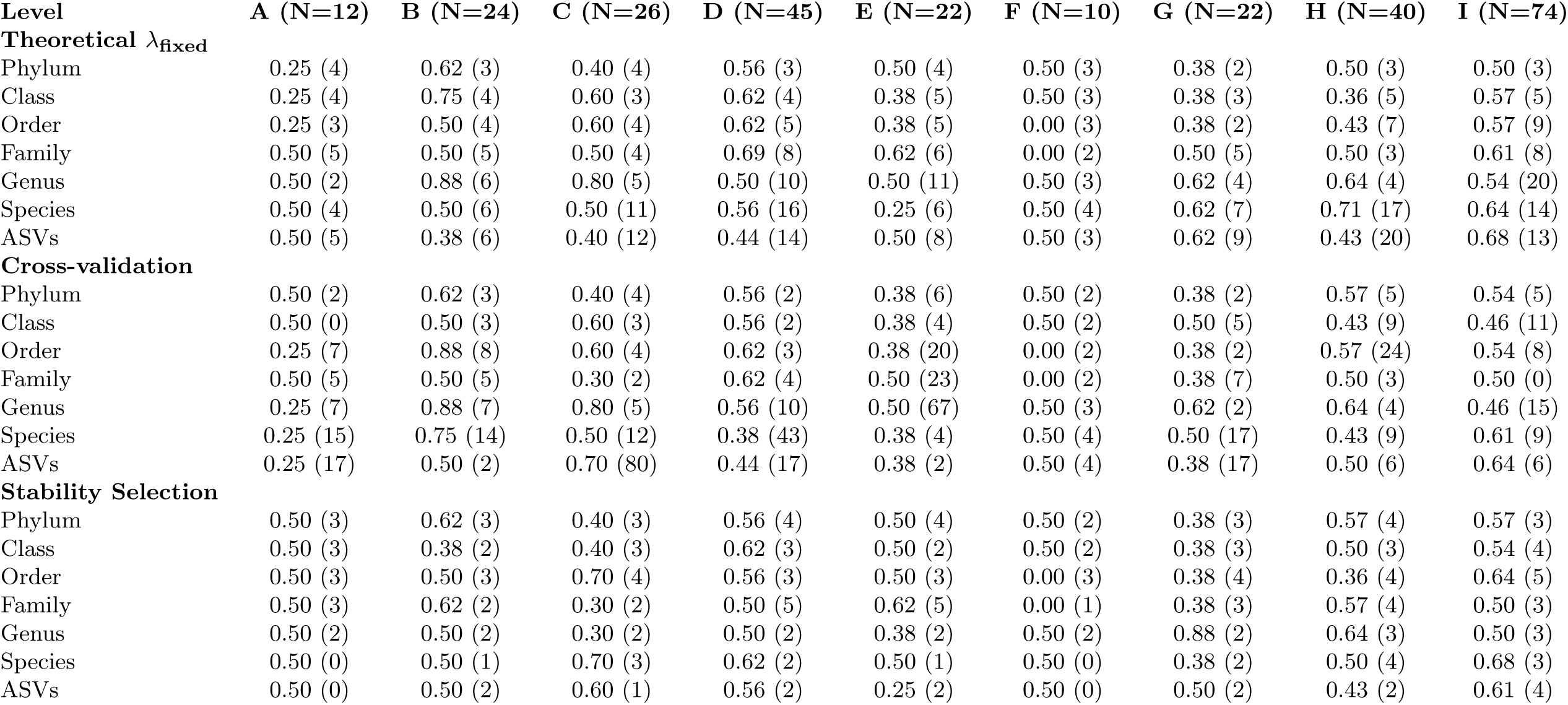
Out-of-sample misclassification rate (MCR) values for IgE clusters (A–I) across taxonomic levels.

| Level | A (N=12) | B (N=24) | C (N=26) | D (N=45) | E (N=22) | F (N=10) | G (N=22) | H (N=40) | I (N=74) |
| --- | --- | --- | --- | --- | --- | --- | --- | --- | --- |
| **Theoretical $\lambda_{\text{fixed}}$** |  |  |  |  |  |  |  |  |  |
| Phylum | 0.25 (4) | 0.62 (3) | 0.40 (4) | 0.56 (3) | 0.50 (4) | 0.50 (3) | 0.38 (2) | 0.50 (3) | 0.50 (3) |
| Class | 0.25 (4) | 0.75 (4) | 0.60 (3) | 0.62 (4) | 0.38 (5) | 0.50 (3) | 0.38 (3) | 0.36 (5) | 0.57 (5) |
| Order | 0.25 (3) | 0.50 (4) | 0.60 (4) | 0.62 (5) | 0.38 (5) | 0.00 (3) | 0.38 (2) | 0.43 (7) | 0.57 (9) |
| Family | 0.50 (5) | 0.50 (5) | 0.50 (4) | 0.69 (8) | 0.62 (6) | 0.00 (2) | 0.50 (5) | 0.50 (3) | 0.61 (8) |
| Genus | 0.50 (2) | 0.88 (6) | 0.80 (5) | 0.50 (10) | 0.50 (11) | 0.50 (3) | 0.62 (4) | 0.64 (4) | 0.54 (20) |
| Species | 0.50 (4) | 0.50 (6) | 0.50 (11) | 0.56 (16) | 0.25 (6) | 0.50 (4) | 0.62 (7) | 0.71 (17) | 0.64 (14) |
| ASVs | 0.50 (5) | 0.38 (6) | 0.40 (12) | 0.44 (14) | 0.50 (8) | 0.50 (3) | 0.62 (9) | 0.43 (20) | 0.68 (13) |
| **Cross-validation** |  |  |  |  |  |  |  |  |  |
| Phylum | 0.50 (2) | 0.62 (3) | 0.40 (4) | 0.56 (2) | 0.38 (6) | 0.50 (2) | 0.38 (2) | 0.57 (5) | 0.54 (5) |
| Class | 0.50 (0) | 0.50 (3) | 0.60 (3) | 0.56 (2) | 0.38 (4) | 0.50 (2) | 0.50 (5) | 0.43 (9) | 0.46 (11) |
| Order | 0.25 (7) | 0.88 (8) | 0.60 (4) | 0.62 (3) | 0.38 (20) | 0.00 (2) | 0.38 (2) | 0.57 (24) | 0.54 (8) |
| Family | 0.50 (5) | 0.50 (5) | 0.30 (2) | 0.62 (4) | 0.50 (23) | 0.00 (2) | 0.38 (7) | 0.50 (3) | 0.50 (0) |
| Genus | 0.25 (7) | 0.88 (7) | 0.80 (5) | 0.56 (10) | 0.50 (67) | 0.50 (3) | 0.62 (2) | 0.64 (4) | 0.46 (15) |
| Species | 0.25 (15) | 0.75 (14) | 0.50 (12) | 0.38 (43) | 0.38 (4) | 0.50 (4) | 0.50 (17) | 0.43 (9) | 0.61 (9) |
| ASVs | 0.25 (17) | 0.50 (2) | 0.70 (80) | 0.44 (17) | 0.38 (2) | 0.50 (4) | 0.38 (17) | 0.50 (6) | 0.64 (6) |
| **Stability Selection** |  |  |  |  |  |  |  |  |  |
| Phylum | 0.50 (3) | 0.62 (3) | 0.40 (3) | 0.56 (4) | 0.50 (4) | 0.50 (2) | 0.38 (3) | 0.57 (4) | 0.57 (3) |
| Class | 0.50 (3) | 0.38 (2) | 0.40 (3) | 0.62 (3) | 0.50 (2) | 0.50 (2) | 0.38 (3) | 0.50 (3) | 0.54 (4) |
| Order | 0.50 (3) | 0.50 (3) | 0.70 (4) | 0.56 (3) | 0.50 (3) | 0.00 (3) | 0.38 (4) | 0.36 (4) | 0.64 (5) |
| Family | 0.50 (3) | 0.62 (2) | 0.30 (2) | 0.50 (5) | 0.62 (5) | 0.00 (1) | 0.38 (3) | 0.57 (4) | 0.50 (3) |
| Genus | 0.50 (2) | 0.50 (2) | 0.30 (2) | 0.50 (2) | 0.38 (2) | 0.50 (2) | 0.88 (2) | 0.64 (3) | 0.50 (3) |
| Species | 0.50 (0) | 0.50 (1) | 0.70 (3) | 0.62 (2) | 0.50 (1) | 0.50 (0) | 0.38 (2) | 0.50 (4) | 0.68 (3) |
| ASVs | 0.50 (0) | 0.50 (2) | 0.60 (1) | 0.56 (2) | 0.25 (2) | 0.50 (0) | 0.50 (2) | 0.43 (2) | 0.61 (4) |

**Table S5.** Summary of alpha diversity analysis results. Sample sizes (N), number of taxa (p), and Mann–Whitney U test p-values for Shannon and Faith’s Phylogenetic Diversity (Faith PD) indices across IgE prevalence groups, IgE burden groups, and IgE clusters.

| Experiment |  | N | p | Shannon (p-value) | Faith PD (p-value) |
| --- | --- | --- | --- | --- | --- |
| IgE Prevalence | Any | 275 | 1441 | 0.5365 | 0.7890 |
|  | Single | 47 | 1075 | 0.2179 | 0.4046 |
|  | Multiple | 228 | 1421 | 0.7827 | 0.5459 |
| IgE Burden | High | 57 | 1084 | 0.8257 | 0.6329 |
|  | Medium | 110 | 1288 | 0.6630 | 0.6171 |
|  | Low | 108 | 1327 | 0.3629 | 0.7601 |
| IgE Clusters | A | 12 | 568 | 0.5857 | 0.6961 |
|  | B | 24 | 829 | 0.9047 | 0.8819 |
|  | C | 26 | 845 | 0.6281 | 0.9614 |
|  | D | 45 | 1050 | 0.5130 | 0.7474 |
|  | E | 22 | 761 | 0.8124 | 0.2100 |
|  | F | 10 | 425 | 0.5336 | 0.0411 |
|  | G | 22 | 773 | 0.8147 | 0.7133 |
|  | H | 40 | 1023 | 0.4181 | 0.9472 |
|  | I | 74 | 1223 | 0.5502 | 0.7391 |

**Table S6.** Taxon Set Enrichment Analysis (TaxSEA) results for the Vitamin A set.

| Set | Statistics | Species |
| --- | --- | --- |
| Vitamin A | Effect Size = -0.013<br>p = $9.88 \times 10^{-1}$<br>FDR = 0.0787 | *Bacteroides massiliensis*, *Dorea formicigenerans*, *Streptococcus salivarius*, *Bacteroides faecis*, *Ruminococcus callidus*, *Parabacteroides goldsteinii*, *Bacteroides salyersiae*, *Bacteroides fluxus*, *Paraprevotella xylaniphila*, *Alistipes indistinctus*, *Acidaminococcus fermentans*, *Clostridium paraputrificum*, *Ruminococcus flavefaciens*, *Streptococcus parasanguinis*, *Lactobacillus mucosae*, *Bacteroides nordii*, *Collinsella tanakaei*, *Odoribacter laneus*, *Megamonas funiformis*, *Streptococcus mutans*, *Parabacteroides gordonii*, *Paraprevotella clara*, *Faecalicoccus pleomorphus*, *Butyricicoccus pullicaecorum*, *Eubacterium limosum*, *Lactococcus garvieae*, *Prevotella corporis*, *Schaalia turicensis*, *Veillonella atypica*, *Porphyromonas uenonis*, *Lactobacillus iners*, *Anaerococcus vaginalis*, *Staphylococcus equorum*, *Haemophilus sputorum* |

**Table S7.**
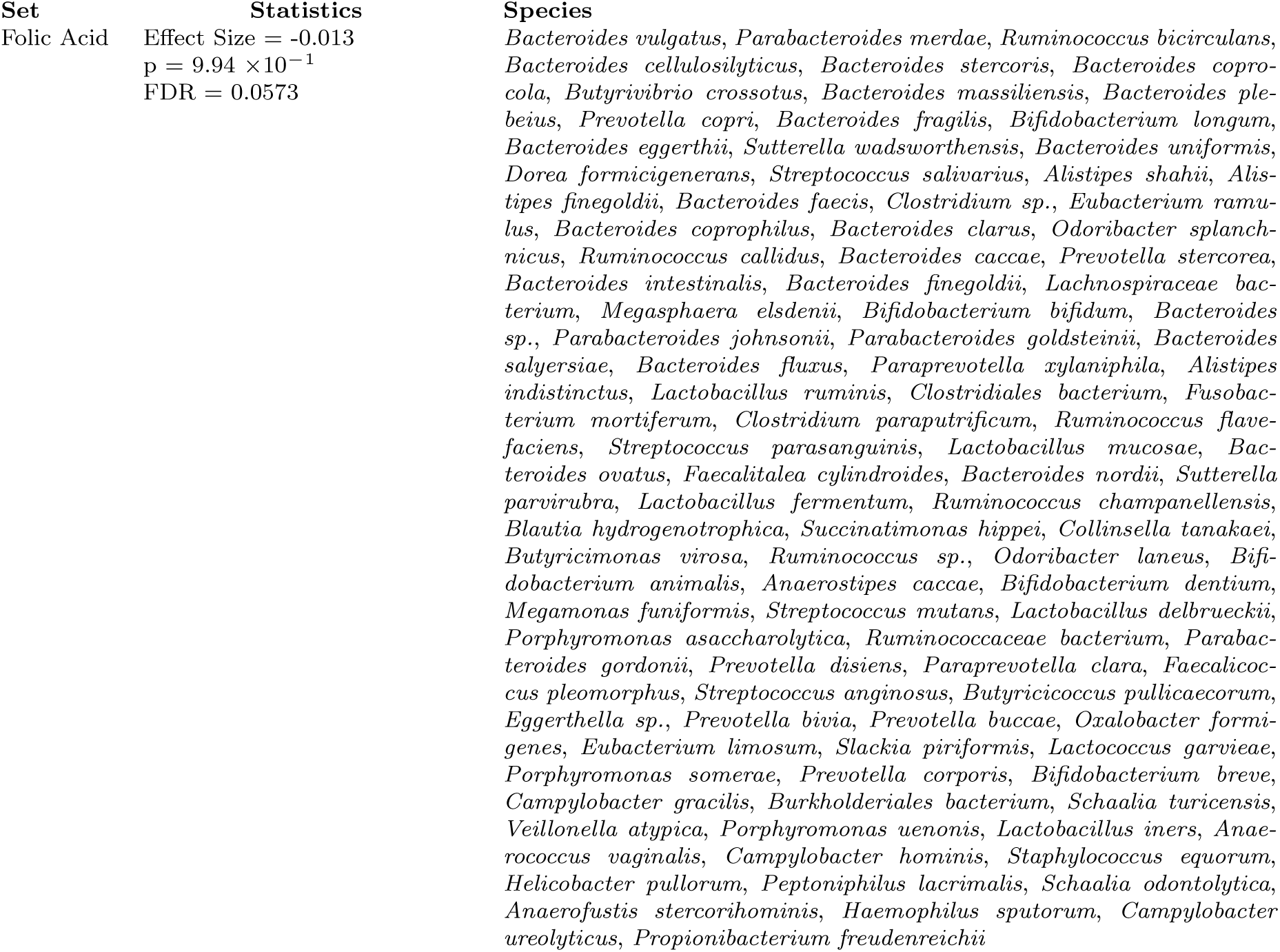
Taxon Set Enrichment Analysis (TaxSEA) results for the Folic Acid set.

**Table S8.**
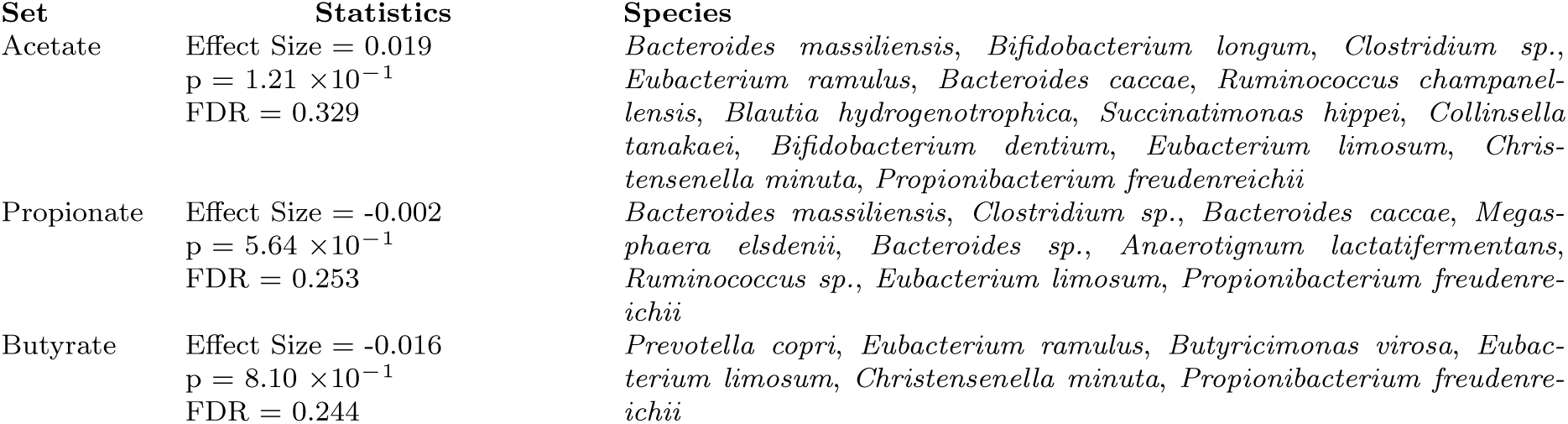
Taxon Set Enrichment Analysis (TaxSEA) results for short-chain fatty acid producer sets.

## STAR METHODS

### Experimental Model and Subject Details

#### Study population

We analyzed data from the KORA FF4 study (2013–2014), the second follow-up of the population-based KORA S4 cohort of residents aged 25–74 years from Augsburg, Germany, and surrounding districts (source population 600,000 in 1999–2001) [34]. Of 6,640 individuals sampled for S4, 4,261 participated at baseline and 2,279 at FF4. For the present study, we included 508 individuals with both allergen-specific IgE measurements and stool microbiome sequencing. Participant characteristics included sex, age, BMI, and physician-diagnosed conditions (hay fever, asthma, dermatitis). All participants provided written informed consent. The study was approved by the Ethics Committee of the Bavarian Chamber of Physicians, Munich (EC No. 06068). A detailed flowchart of participant selection and data integration is shown in Figure S1.

### Method Details

#### High-Throughput 16S rRNA Gene Amplicon Sequencing

Stool samples were collected using tubes containing 5 ml stool stabilizer (Invitek DNA Stool Stabilizer, No. 1038111100). DNA was extracted following [35]. Amplicon libraries targeting the V3–V4 region were generated with primers 341F (CCTACGGGNGGCWGCAG) and 785R (GACTACHVGGGTATCTAATCC) [36]. PCR was carried out using Phusion High-Fidelity DNA Polymerase HotStart (Thermo Fisher, Cat# F-540S), and products were barcoded with dual-index primers, purified using AMPure XP beads (Beckman Coulter), quantified, and pooled at 2 nM. Sequencing was performed on an Illumina MiSeq platform (2 x 250 bp) at the ZIEL NGS-Core Facility, Technical University of Munich.

#### IgE screening

Allergen-specific IgE antibodies were measured using the ImmunoCAP ISAC 112 microarray platform (Thermo Fisher Scientific) according to the manufacturer’s protocol. This platform covers 112 allergen components spanning food, pollen, house dust mite, and animal dander. Sensitization was defined by the presence of detectable IgE (*≥*0.3 ISU), without applying the conventional 0.35 kUA/L threshold, in order to include individuals with low-level responses. IgE-positive individuals were selected based on survey-reported allergy, while a subset of non-allergic controls was randomly chosen for comparison. A full list of allergen components is provided in Supplementary Tables.

#### Bioinformatic processing

Raw reads were analyzed using QIIME2 v2021.11 [37]. Demultiplexing and denoising were performed with q2-demux and q2-dada2 [38]. Taxonomic assignment of ASVs [39] was performed with q2-feature-classifier [40] using the sklearn naïve Bayes classifier [41] trained on the SILVA 138 SSU database [42]. ASVs present in *≤*1% individual were removed, resulting in 6,958 ASVs for downstream analysis.

### Quantification and statistical analysis

IgE co-sensitization patterns were characterized using hierarchical clustering and singular value decomposition (SVD), followed by Varimax rotation to define latent allergy components (LACs) [19]. IgE co-occurrence networks were constructed by computing co-occurrence frequencies across the entire study population. Microbial alpha diversity was assessed using Shannon and Faith’s Phylogenetic Diversity indices, compared between groups using Wilcoxon rank-sum tests with Benjamini-Hochberg correction at level 0.1 [43]. Differential abundance analysis was conducted with LinDA v1.0 [20] at the ASV, species, and genus levels. Presence/absence analyses were performed to identify group-specific taxa. Classification was performed using sparse log-contrast classification with squared hinge loss using the c-lasso Python package[24]. For sparse model selection, we used the three built-in options: cross-validation [44], stability selection [45], and a fixed theoretically-derived, asymptotically optimal regularization strength [25, 26]. Family-level microbial association networks were inferred using SPIEC-EASI [29] with graphical lasso and StARS stability selection criterion [46] via PULSAR [47] for regularization parameter selection, applied to the set of all microbial families. Networks were constructed and compared between IgE groups using the NetCoMi R package [30], with differential networks identifying associations that were gained, lost, or reversed. Taxon Set Enrichment Analysis (TaxSEA) [31] was applied to identify enriched functional groups, using reference sets from the Human Microbial Metabolome Database [48].

